# Targeted inCITE-Seq Analysis Identifies the Loss of Nuclear TDP-43 in Endothelium as a Mediator of Blood Brain Barrier Signaling Pathway Dysfunction in Neurodegeneration

**DOI:** 10.1101/2023.12.13.571178

**Authors:** Omar M.F. Omar, Amy L. Kimble, Ashok Cheemala, Jordan D. Tyburski, Swati Pandey, Qian Wu, Bo Reese, Evan R. Jellison, Yunfeng Li, Bing Hao, Riqiang Yan, Patrick A. Murphy

## Abstract

Despite the importance of the endothelium in the regulation of the blood brain barrier (BBB) in aging and neurodegenerative disease, difficulties in extracting endothelial cell (EC) nuclei have limited analysis of these cells. In addition, nearly all Amyotrophic Lateral Sclerosis (ALS) and Frontotemporal Degeneration (FTD), and a large portion of Alzheimer’s Disease (AD) exhibit neuronal TDP-43 aggregation, leading to loss of nuclear function, but whether TDP-43 is similarly altered in human BBB ECs is unknown. Here we utilize a novel technique for the enrichment of endothelial and microglial nuclei from human cortical brain tissues, combined with inCITE-seq, to analyze nuclear proteins and RNA transcripts in a large cohort of healthy and diseased donors. Our findings reveal a unique transcriptional signature in nearly half of the capillary endothelial cells across neurodegenerative states, characterized by reduced levels of nuclear β-Catenin and canonical downstream genes, and an increase in TNF/NF-kB target genes. We demonstrate that this does not correlate with increased nuclear p65/NF-kB, but rather a specific loss of nuclear TDP-43 in these disease associated ECs. Comparative analysis in animal models with targeted disruption of TDP-43 shows that this is sufficient to drive these transcriptional alterations. This work reveals that TDP-43 is a critical governor of the transcriptional output from nuclear p65/NF-kB, which has paradoxical roles in barrier maintenance and also barrier compromising inflammatory responses, and suggests that disease specific loss in ECs contributes to BBB defects observed in the progression of AD, ALS and FTD.

## Introduction

The blood brain barrier (BBB) tightly regulates the molecular and cellular composition of the central nervous system ^1^. Barrier properties are derived from the function of the tight junction proteins such as Cldn5 and Occludin, which limit passive transfer of molecules and proteins across the endothelium, and is developmentally induced by Wnt signaling activity ^2^. Endothelial cell functions are supported by a basement membrane and adjacent cells, including pericytes, astrocytes, perivascular fibroblasts and smooth muscle cells, but the essential barrier functions are intrinsic to the endothelium. Alterations in transcellular transport and paracellular diffusion of molecules have been observed in aging, as paracellular leak increases ^3,4^ and transcellular transport is reduced ^5^. These alterations in the aging brain may not be benign, as larger changes in paracellular leak are observed at the earliest stages of Alzheimer’s Disease (AD), and are a harbinger of cognitive decline ^6,7^.

The cellular mechanisms of BBB dysfunction in AD are not yet fully clear, but the observation of similar dysfunction in ALS and FTD suggest the possibility that the diseases may share some underlying mechanisms. Unlike AD, ALS and FTD are relatively early onset, and driven in large part by genetic lesions which directly or indirectly affect the nuclear localization of the RNA-binding protein TDP-43 (*TARDBP*). TDP-43 is reduced in the neuronal nuclei nearly all cases of ALS and approximately half of the cases of FTD ^8^. Many AD cases exhibit a very similar pattern of neuronal TDP-43 aggregation to FTD cases, suggesting that similar underlying mechanisms may be involved ^9^. Notably, TDP-43 dysfunction is not confined to neurons, and is observed in astrocytes ^10,11^ in the brain, the pancreatic islet ^12^, and even fibroblasts isolated from skin ^13^. Although the level of expression of TDP-43 is relatively high within the endothelium, and comparable to neurons and fibroblasts, changes in the nuclear levels of TDP-43 in the endothelium in aging and disease have not been carefully examined.

Despite an increasing amount of single cell data from human tissue samples, several important hurdles remain in understanding the alterations of the BBB in aging and neurodegenerative disease. First, analysis of endothelial cells in donor derived samples has been hampered by the low recovery in single nuclei preparations ^14,15^. Methods to enrich for brain microvasculature specifically have yielded important insight, but require substantial processing steps, and have not yet been applied to large sets of samples ^16^. In addition, a focus on transcriptomic techniques has limited an understanding of alterations in proteins, including splice factors and transcription factors, where function is determined not by transcript level, but by nuclear levels and activity – like TDP-43. Other key transcription factors involved in BBB function, including RELA/p65 (NF-kB signaling) CTNNB/β-catenin (Wnt signaling) are also uncharted in alterations with age and in neurodegeneration. So, despite substantial information on these pathways in animal models, the relevance to age and disease associated changes in the human BBB remain unknown.

Here, we apply a technique we recently developed for the specific enrichment of endothelial and microglial nuclei from frozen brain tissues^17^ together with <u>in</u>tranuclear <u>C</u>ellular <u>I</u>ndexing of <u>T</u>ranscriptomes and <u>E</u>pitopes (inCITE-seq), a method for the analysis of nuclear proteins together with RNA transcripts, to a large cohort of human cortical brain tissues from healthy donors, young and old, and donors with ALS, FTD and AD. Surprisingly, we found a tight correlation between nuclear levels of TDP-43 and p65/NF-kB in healthy endothelial cells which was disrupted in all disease conditions associated with neurodegeneration. In these TDP-43 deficient cells, NF-kB was associated with a strong transcriptional response not observed in unaffected endothelial cells, and the Wnt signaling pathway critical for BBB integrity was impaired. These defects were recapitulated in brain endothelial cells *in vitro* and *in vivo* with targeted loss of TDP-43, implicating a reduction in nuclear TDP-43 levels in both alterations in neurodegenerative endothelium.

## Materials and Methods

### Antibody-Oligonucleotide conjugation

The protocol was followed as outlined previously, with some modifications (https://citeseq.files.wordpress.com/2019/03/cite-seq_hyper_conjugation_190321.pdf). TCO-PEG4-Oligo Labeling was performed according to the original protocol without any changes. The purchased antibodies (refer to Table 2), which were generally BSA- and azide-free. If antibodies included BSA as a stabilizer, they were cleaned with Melon columns (Thermo). Antibodies cleaned twice using a pre-wet Amicon Ultra-0.5 30 kDa MWCO filter, as suggested in the protocol. Subsequently, the columns were inverted, and 5 µl of diluted 4mM mTz-PEG4-NHS was added, followed by a 30-minute incubation for antibody functionalization. To quench the reaction, 1 µl of 1 M glycine (pH 8.5) was added. Any excess mTz-PEG4-NHS was removed by filtering through the same 30 kDa MWCO filter for 5 minutes at 14,000g. The filter was then inverted into a clean collection tube and spun at 100g for 2 minutes to elute the product. The functionalized antibody was subsequently allowed to react with TCO-PEG4-oligo. For every 1 µg of antibody, 30 pmol of TCO-PEG4-oligo was added. The reaction mixture was then incubated overnight in a 4°C fridge. After the reaction was complete, 1/10th of the reaction volume of 10 mM TCO-PEG4-glycine was added to quench any residual tetrazine reaction sites on the antibody. Conjugation was verified using western blot analysis, which checked for an increase in the size of the heavy chain after conjugation and laddering for the hyperconjugated product (SI Figure 8). To eliminate excess oligonucleotides, the reaction mixtures of individual antibody/oligonucleotide pairs were combined and subjected to multiple ammonium sulfate precipitation steps for purification. The resulting elutes containing antibodies were applied to a size exclusion chromatography column (Superdex 200 10/300 GL; GE Healthcare) with PBS served as running buffer and a flow rate of 0.5 mL/min (SI Figure 8). The elution of the factions was monitored simultaneously at wavelengths 280 nm and 260 nm using an AKTA pure chromatography system equipped with a UV monitor U9-M (SI Figure 8).

### Enrichment of endothelial nuclei from frozen post-mortem brain tissue

Nuclei isolation generally followed our previously published protocol with minor modifications ^17^. All steps were performed either on ice or at a temperature of 4°C, unless specified otherwise. Frozen cortical brain tissues, weighing between 100 to 200 mg, were thawed at room temperature for 3-4 minutes in Nuclei EZ lysis buffer (Sigma Nuc 101) mixed with RNAase inhibitor (0.5U/µl, Clontech Cat #2313) in RINO tubes containing eight 3.2 mm stainless steel beads. The tissues were then mechanically homogenized using a Next Advance Bullet Blender BB724M, set to level 4 for 4 minutes at 4°C. Following homogenization, the mixture was diluted in 5 mL, centrifuged at 500xG for 5 minutes (retaining the supernatant for analysis of total brain protein and mRNA), and subsequently washed with 5 mL of Nuclei EZ lysis buffer. A 2-minute incubation on ice preceded a second centrifugation at the same speed for another 5 minutes. The supernatant was then discarded, and the homogenate was resuspended in 5 mL of Nuclei EZ lysis buffer, followed by a 5-minute incubation on ice. The homogenate was then strained through a 70 µm pluristrainer, centrifuged at 4°C, and the supernatant removed. The resulting nuclei pellet was suspended in 200 µl of a PBS-based staining solution (1% BSA, PBS, 0.5U/µl RNAase inhibitor) in 1.5 ml low-bind tubes. After washing the nuclei with staining buffer (PBS + 0.2% BSA + RNAse inhibitor), the supernatant was removed, and 50 µl of blocking buffer, containing ssDNA (1 mg/mL final concentration) and FC block (1:100 of BioLegend, 156604), was added. This was followed by a 10-minute incubation. The nuclei were then stained with a buffer containing Anti-Erg 647 (1:200, Clone EPR3864, Abcam), Anti-NeuN Cy3 (1:200, Sigma), DAPI for nuclei labeling (1:2000 of a 5 mg/mL stock), and InCITE seq antibody mix. Before using the antibody mix, we incubated it with EcoSSB (Promega M3011) in 50 μl of 1X NEBuffer 4 for 30 min at 37 °C as suggested previously ^18^. Afterwards, 1 µl of TotalSeqB Hashtag (anti-Nuclear Pore Complex Proteins Hashtag at 0.5g, Biolegend 682239-682243) was added to each sample for subsequent identification in downstream analysis. After staining, the nuclei were washed twice with staining buffer and fixed with 2% PFA in staining buffer for 1 minute at 4C. Then, 500 µl of staining buffer containing 0.1% glycine was added to quench the fixation. The nuclei were centrifuged for 5 minutes at 500xG twice and resuspended in 500 µl of staining solution. This solution was then filtered through a 35 µm filter before sorting the nuclei using an Aria 2 with a 40 µm nozzle, collecting them into a BSA-coated tube maintained at 4°C.

### Droplet based 10x genomics single nucleus RNA sequencing

For human tissues, nuclei from 4 to 5 human donors were pooled to a final volume of 42 µl. We aimed for each 10x reaction to include two control groups (one from an older healthy individual and another from a younger one) and two groups representing different disease states (AD, ALS, or FTD), each uniquely tagged with a nuclear pore hashtag for subsequent identification. We aimed for total nuclei count in each mixture to be ∼8000 nuclei. These mixtures were then loaded onto a Chromium single-cell V3 3′ chip (10x Genomics) and processed according to the manufacturer’s protocol. Briefly, after GEM generation, they were subjected to reverse cross-linking and reverse transcription by heating at 53°C for 45 minutes and then at 85°C for 5 minutes. The samples were subsequently stored at −20°C until GEM recovery. For cDNA amplification, the standard 10x Genomics protocol was used, incorporating totalseqB PCR additive primers into the mix. Following amplification, a SPRI-based size selection separated antibody-oligonucleotide-derived cDNA fragments from mRNA-derived cDNA. The antibody-oligonucleotide cDNA, contained in the supernatant, was set aside for antibody capture library construction. Meanwhile, the mRNA-derived cDNA, attached to SPRI beads, was processed for gene expression library construction. Gene expression libraries were prepared from the mRNA-derived cDNA using enzymatic fragmentation, adaptor ligation, and sample indexing, following the 10x Genomics protocol. These libraries were stored at −20°C until quantification and sequencing. For antibody capture library, the antibody-oligonucleotide-derived cDNA underwent mixing with 1.4× SPRIselect, magnetic separation, and ethanol washes, followed by elution in sterile water. PCR amplification of this cDNA, using unique TotalSeq-B index primer mixes, created the protein capture libraries. These libraries were then purified, eluted, and stored at 4°C until quantification and sequencing on a NovaSeq 6000 using version 1.5 chemistry.

### inCITE-seq and snRNA-seq data preprocessing

Sequencing data were processed with Cell Ranger version 7.0.1 on Xanadu high performance computing cluster. Reads from FASTQ files were aligned to mouse mm10 or human GRCh38 reference genome as described by 10x genomics. Paths to feature library (antibody capture and gene expression) were specified during the processing, so we ended with a h5 object that included both gene expression and antibody capture information. Hashed nuclei were demultiplexed using DemuxEM using Pegasus ^19^ implementation in Python2 with parameters min_signal=15.0, alpha=0.0, alpha_noise=150.0, and deMULTIplex2 ^20^. nuclei with ambiguous hashtag assignment were discarded. All downstream analysis were performed using the Scanpy (Version 1.8.1) ^21^. One expected sample was not found in batch Jan19B_2023 (NuclearPoreHash2) by hashtagdemultiplexing.

### snRNA sequencing analysis

Nuclei were required to express at least 50 genes, and each gene had to be present in a minimum of 5 cells. Nuclei were excluded if they exhibited more than 5% mitochondrial gene content, had Nuclear pore hashtag counts exceeding 4000, or displayed anti-TDP43 antibody levels above 3000 Antibody counts counts. Adhering to these parameters, the dataset was narrowed down to 149862 nuclei and 32,767 genes. The normalization process for gene counts in each nucleus involved individual normalization followed by logarithmic scaling using the formula ln(x + 1).

Cell type clustering (Figure 1) : For the clustering of cell types, 6,830 variable genes were identified using the ’highly_variable_genes’ function in Scanpy, with the parameters set to min_mean=0.0015, max_mean=0.18, and min_disp=0.30. Log counts were scaled, and UMI counts were adjusted using the ’regress_out’ function in Scanpy. Dimensionality reduction was then carried out on these variable genes using PCA in Scanpy, employing the arpack solver. To account for batch effects, Harmony was applied ^22^, focusing on corrections for 10x well batches and individual human samples effects. Then, a nearest-neighbor graph was constructed using the top 40 principal components and k=10 neighbors, followed by clustering using the Leiden algorithm at a resolution of 0.8. The data was then visualized using UMAP. Further 9169 suspected doublets were removed using scrublet, ending with 140693 nuclei. After removal of those nuclei, PCA and harmony corrected PCA were recalculated to account for reduced number of nuclei. Celltypes were then annotated using Pegasus annotate function with nuclei with indeterminate assignment or could be identified by looking their top genes were removed or high in mitochondrial counts were removed ending up with 139342 cells.

**Figure 1.**
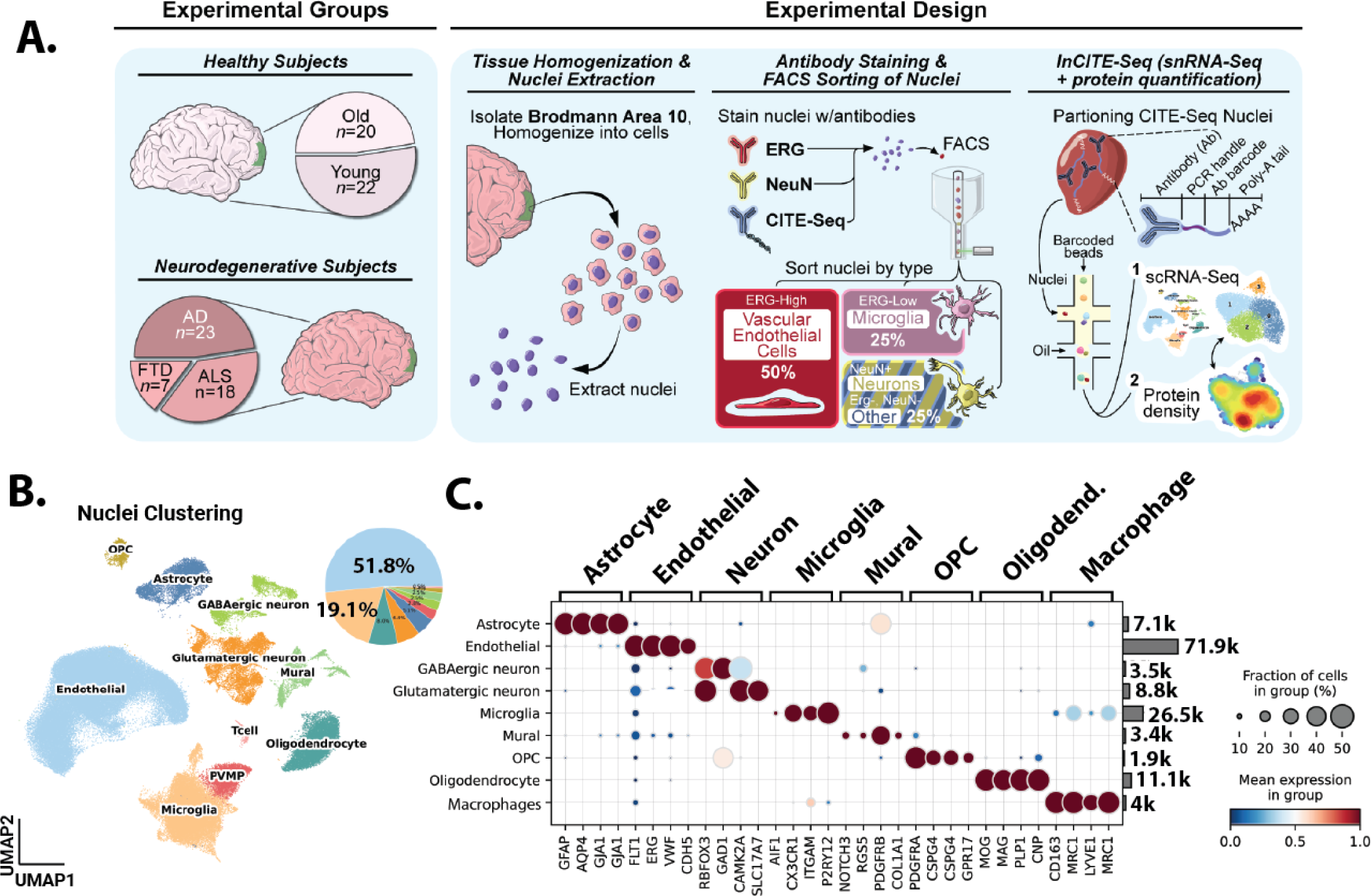
Single nuclei analysis of human tissue samples with endothelial and microglial enrichment. (A) Schematic representation of the methodology employed for endothelial enrichment, leveraging ERG and intranuclear CITEseq antibodies. (B) Uniform Manifold Approximation and Projection (UMAP) visualizing ∼140,000 nuclei from 89 human frontal cortex samples, color-coded by cell type, and pie chart depicting the distribution of captured cell types. (C) A dual representation featuring a dot plot of cell type-specific gene markers and a bar chart illustrating the count of captured nuclei for each cell type.

**Figure 2:**
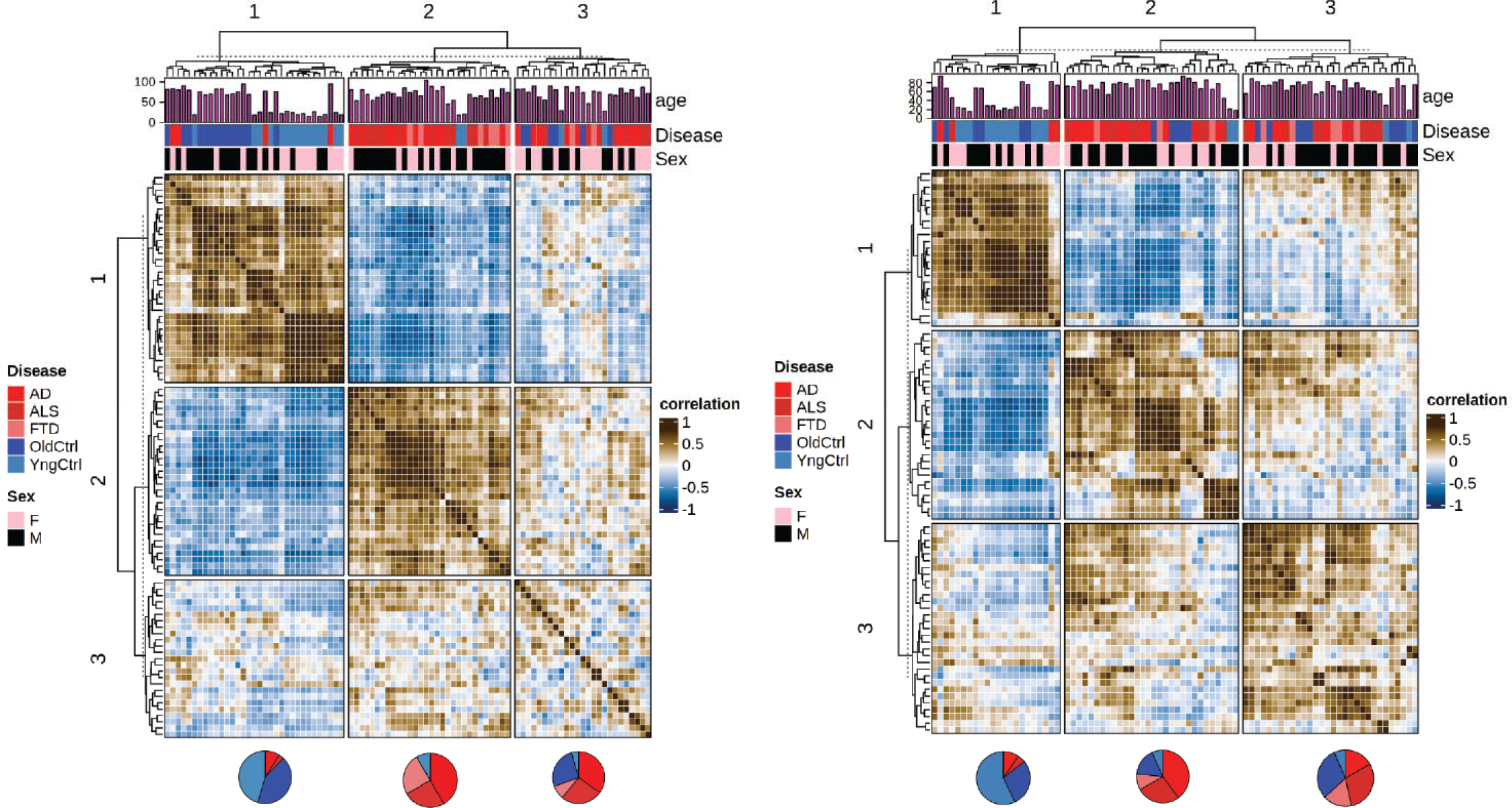
Principal component analysis reveals shared transcriptional processes across neurodegeneration. (A) A correlation presents Pearson correlations of PCA-adjusted capillary endothelial cell profiles from diverse donors, processed via Harmony for integration based on batches. The PCA, based on 50 components, is followed by hierarchical clustering and dendrogram visualization, categorizing data by sex and disease state. (B) Similar correlation matrix for microglial population.

Disease specific clustering (Figures 3-6): As clustering to define cell types used Harmony to limit variation due to disease state, we aimed to return to major cell type classifications derived by original clustering to specifically examine similarities in transcriptional states between disease states and donors. We focused on two major groups, capillary endothelial cells and microglia. For each, nuclei associated with this cluster were isolated in silico, and then re-clustered, this time regressing out only effects of the 10x well batch, and not disease state or donor.

**Figure 3:**
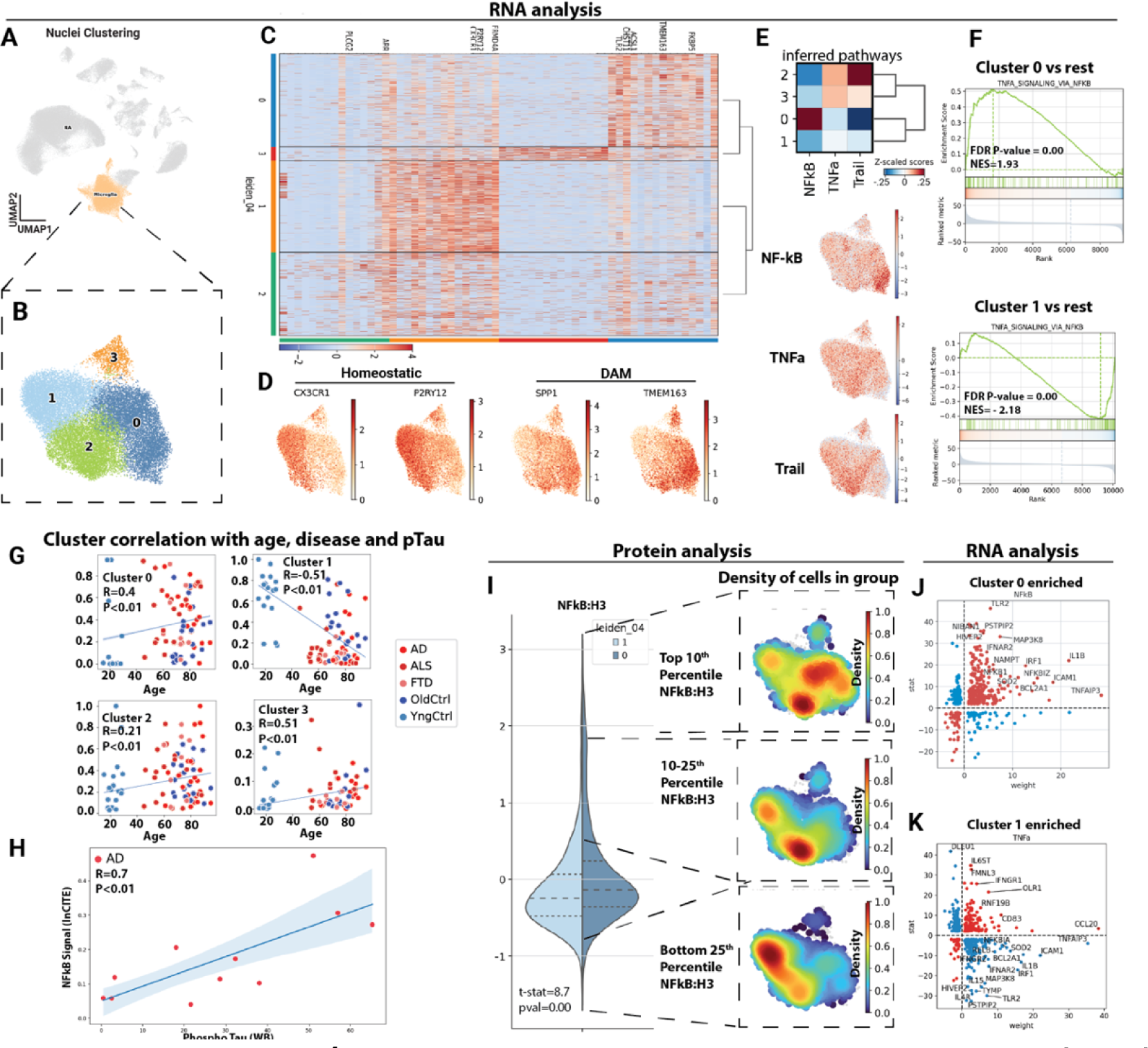
Microglial p65/NFkB associates with disease associated microglia (DAM). (A) Uniform manifold approximation and projection (UMAP) clustering representation of cortical microglial cells, derived from ∼ 39K cells in the inCITE dataset, with disease state regressed out. (B) *in silico* isolation of microglia nuclei and uniform manifold approximation and projection (UMAP) clustering based on batch corrected gene expression patterns, without regression of disease state. (C) Heatmap illustrating differential gene expression patterns across the identified microglial subclusters. (D) Gene counts for disease associated microglia (DAM) and homeostatic markers across cells. (E) Comprehensive analysis of inferred pathways prevalent in the microglial clusters by Progeny and mapping of inferred pathway scores onto cells in UMAP, red signal indicates higher expression of pathway genes. (F) GSEA analysis of TNF-alpha signaling pathway within clusters 0 and 1 in contrast to other clusters from MsigDB. (G) Scatter plot showing correlations (Spearmann) between donor age and disease status and its respective contribution to individual microglial clusters and (H) scatter plot showing correlation between the number of nuclei in DAM-enriched cluster 0 and pTau levels determined by Western blot in the same brain tissue. (I) Violin plot showing the mean level of p65/NFkB protein (normalized to levels of histone H3 protein) in cells of microglial cluster 1 and 0, and informatic isolation of nuclei with highest (Top 10^th^ percentile), high (Top 10-25^th^ percentile) and low (Bottom 25^th^ percentile) levels of NFkB:H3 shown as a density plot. Areas of the UMAP enriched for cells falling into this range of NFkB:H3 are red, lower levels are blue. (J) Scatter plot showing gene weight (relative association in the NFkB pathway as predicted by the Progeny database, X-axis) and log fold change between Cluster 0 and other clusters (Y-axis). In red are transcripts increased or decreased in a manner consistent with NFkB activation, and in blue are transcripts following a pattern of reduced NFkB activity. (K) Similar plot for Cluster 1.

**Figure 4:**
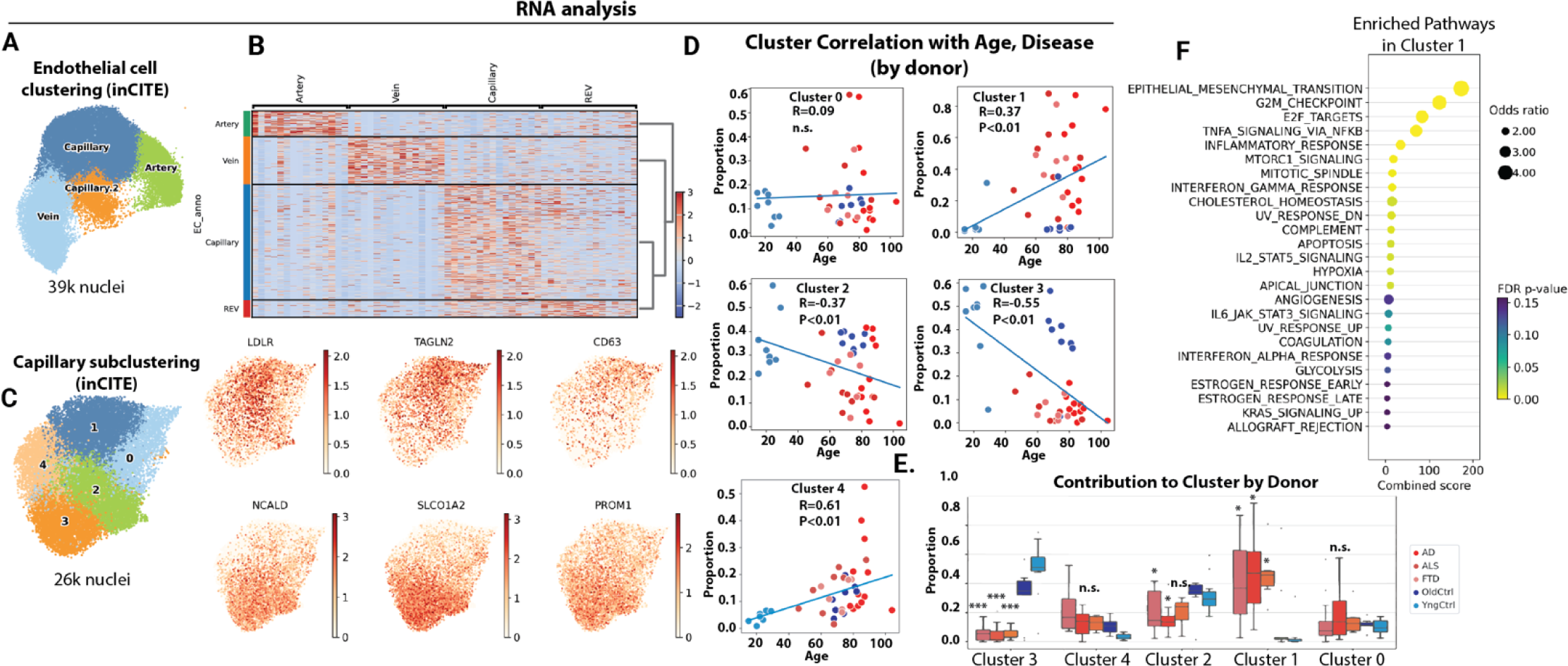
Distinct brain capillary endothelial states associate with healthy aging versus neurodegenerative diseases. (A) Uniform manifold approximation and projection (UMAP) clustering representation of brain endothelial cell subtypes, encompassing capillaries, veins, and arteries, derived from ∼39K cells in the inCITE dataset, with disease state regressed out. (B) Heatmap showing identification markers for capillary endothelial cell subtypes. (C) *in silico* isolation of capillary nuclei and UMAP clustering based on batch corrected gene expression patterns and without regression of disease state. Data represents ∼26K cells and feature plots of gene expression for cluster marker genes enriched in cluster 1 or cluster 3 relative to others. (D) Scatter plot showing correlation (Spearmann) between donor age and contribution to each capillary cluster, and (E) box plot showing the proportions of capillary endothelial cells for each donor falling into each cluster. (F) Bubble plot showing pathway enrichment from MsigDB in disease associated cluster 1 versus all other clusters. (E) ANOVA with post-hoc Tukey test, ***P<0.001, *P<0.05.

**Figure 5.**
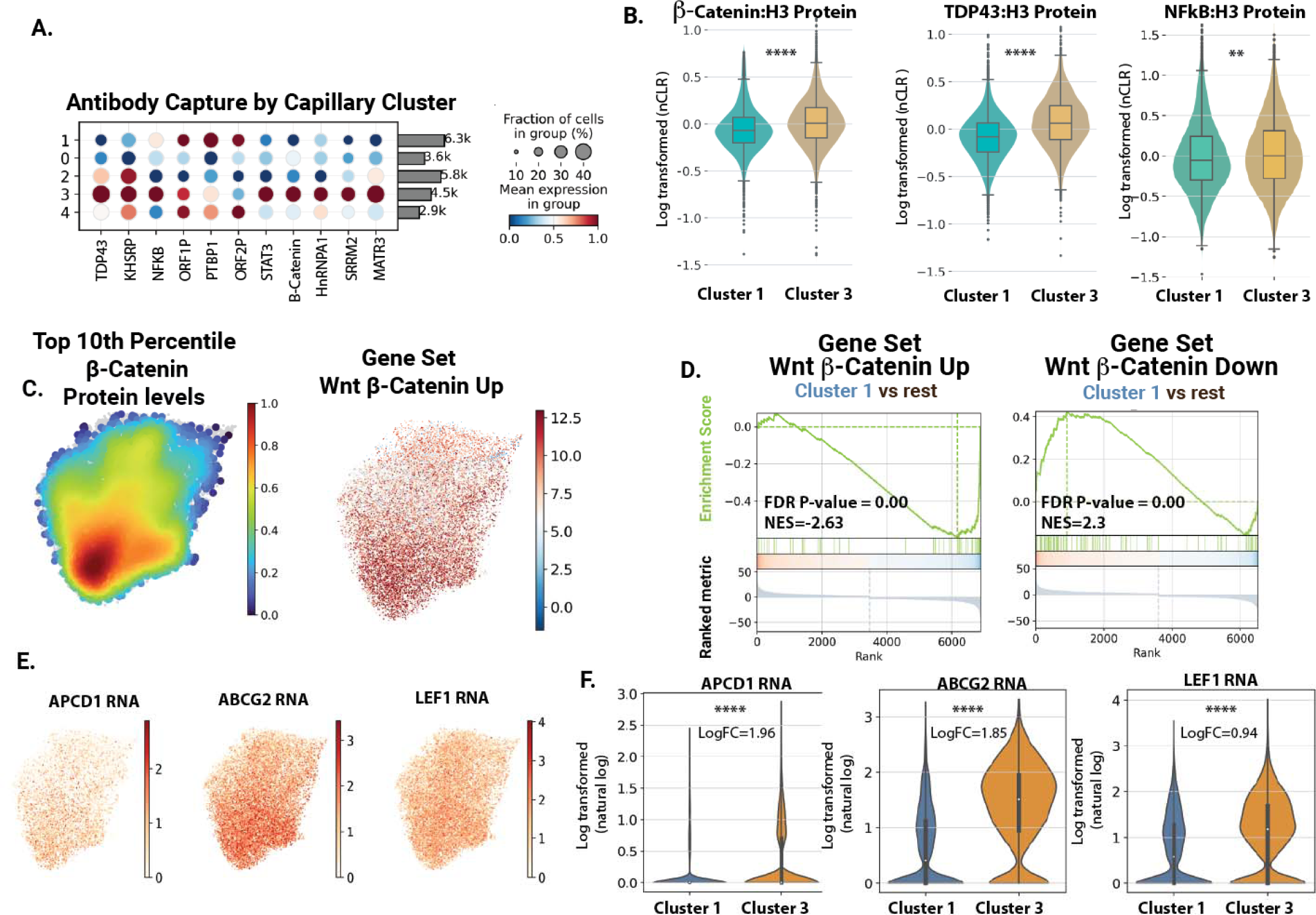
inCITE analysis reveals a loss of Wnt/β-Catenin signaling in disease associated nuclei. (A) Dot plots showing relative levels of protein across all cell clusters, and within the endothelial and microglial clusters. (B) Violin plot showing histone normalized protein levels for p65/NF-kB, TDP-43 and Beta-Catenin in Cluster 1 (disease cluster) in comparison to Cluster 3 (healthy control). (C) Density plots illustrating the top 10th percentile of protein levels for Beta-Catenin (relative to H3), and predicted positive regulation of Wnt signaling pathway by gene expression. (D) GSEA plot of Wnt/β-Catenin genes enriched in disease associated cluster 1 (versus all other clusters). Gene sets are either genes positively associated with Wnt activation in human endothelial cells, or negatively associated with Wnt activation. (E) Feature plots showing example genes in the β-Catenin pathway, projected onto the capillary UMAP. (F) Violin plots showing Wnt target gene expression across clusters.

**Figure 6.**
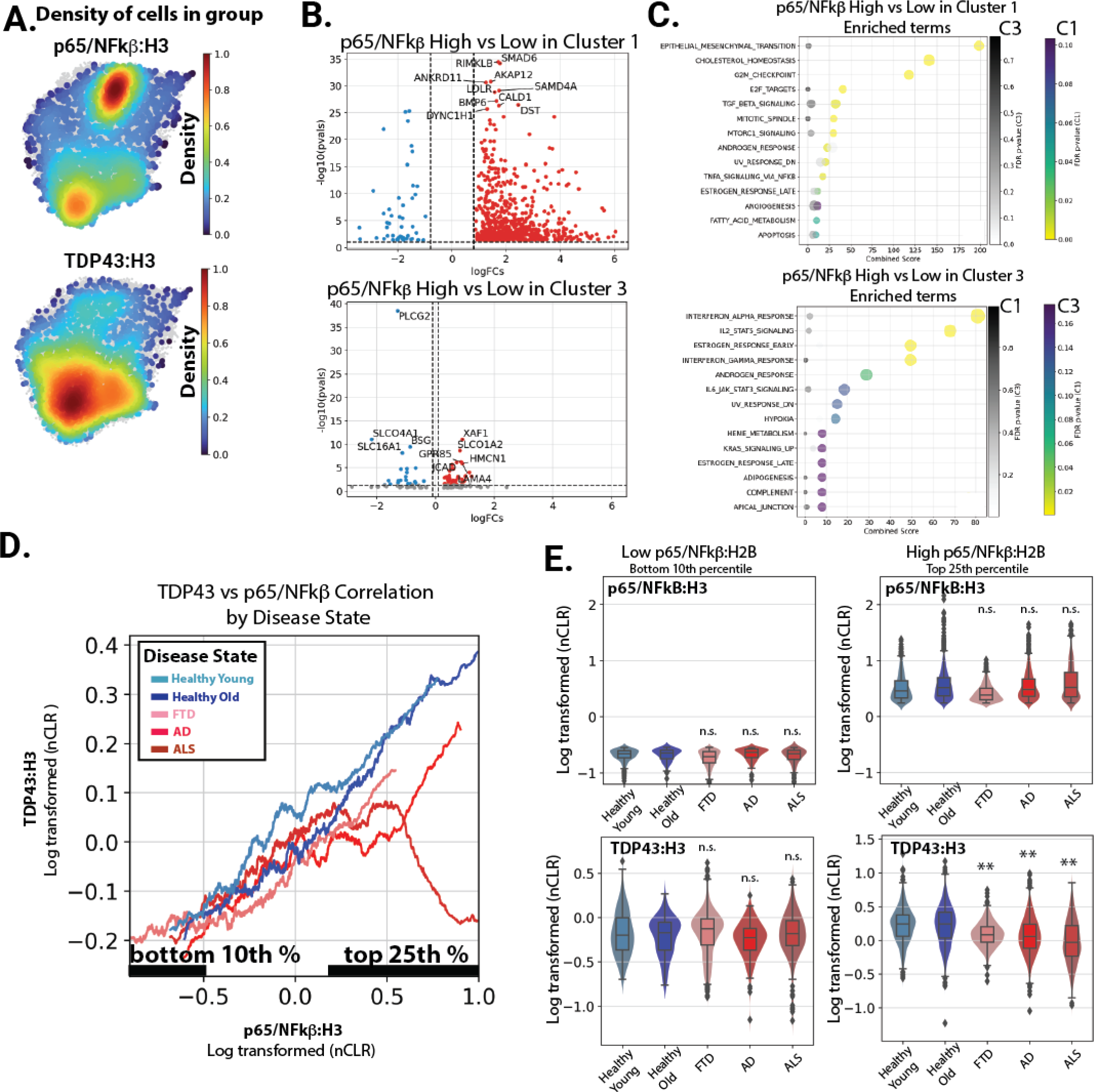
inCITE-seq analysis reveals a specific loss of nuclear TDP-43 and increased NFkB transcriptional targets disease associated nuclei. (A) Density plots illustrating the top 10^th^ percentiles of protein levels for p65/NFkB, Beta-Catenin, and TDP-43. (B) Transcripts positively associated with the highest (top 10^th^ percentile) of nuclear p65/NFkB relative to the lowest (bottom 25^th^ percentile) in each cluster. (C) GSEA Enrichr analysis of MSigDB pathways highest in the genes upregulated with p65/NFkB in cluster 1 and cluster 3, respectively. Enrichment of the pathways shown for each cluster are also shown in grey for the other cluster. (D) A smoothed line plot comparing NFkB protein levels (x-axis) to TDP-43 (y-axis). (E) Violin plot showing histone normalized protein levels for p65/NFkB and TDP-43 in bottom 10^th^ percentile and top 25^th^ percentile of p65/NFkB. Significance relative to healthy aged control is shown.

### Protein Capture normalization

We normalized the protein amount as previously reported, except we normalized to H3 histone, instead of nuclear pore protein hash ^18^. This change was made because the nuclear pore protein hash showed greater variability than H3 histone across different disease states and nuclei (SI Figure 5). This normalization was achieved using the formula: n(Antibody capture counts) = (Antibody capture counts) / Histone H3 Counts. Following this, we transformed these normalized values into centered natural log ratios, termed ’nCLR’, using the formula: nCLR = n(protein_capture)/ (∏in(protein_capture)i ^(1/n). Here, the denominator represents the geometric mean, calculated by taking the nth root of the product (∏) across each nucleus (i). The normalized protein data were visualized using density plots, which were representative of the quintiles of protein expression. We predefined these quintiles and labeled each individual cell observation with the quintile range it fell into. For further analysis, we employed the embedding_density function from the Scanpy tools suite. This function was used to calculate an embedding density protein, which was then visualized using the embedding_density plotting function.

### Pathways and gene set enrichment analysis

We identified marker genes and differentially expressed genes between clusters using the t-test method, as implemented in the rank_genes_groups() function in Scanpy. For microglia pathway analysis, we utilized the PROGENy database implemented in Python ^23^. Specifically, we exported the top 1000 terms from the PROGENy database. We then compared the inferred Leiden cluster pathways with inferred cellular activity on a per-cell basis using the run_mlm function. The results were saved in the obsm of the Anndata object and visualized through a heatmap and on a UMAP to highlight differences in specific inferred pathways. Genes downstream of the pathways in PROGENy were visualized using a scatter plot, plotting the weight from the PROGENy table on the x-axis against log fold changes from the differential gene expression analysis. Enrichment and Gene Set Enrichment Analysis (GSEA) were performed using the exported differential gene expression table. For NF-kB-related pathways, we utilized the curated Hallmark gene sets from MsigDB. For β-catenin analysis, we employed custom-curated gene sets derived from studies on β-catenin function modulation in pluripotent stem cell-derived endothelial cells ^24^. Similar gene set signatures were generated from other datasets generated in the lab or publicly available (SI Table 3). In our cell proportion analysis, we assessed the relative contribution of disease states in clusters to identify common associations in gene expression (GEX) in dementia. This analysis involved visualizing the relationship between age and the proportion of cells in each cluster using scatter plots with regression lines. We quantified this relationship using Spearman’s rank correlation.

### Western analysis of protein supernatant from nuclei isolation

After centrifugation to spin nuclei down from EZ-lysis buffer, samples were stored at −80C. Protein quantification was performed by BCA assay (Thermo 23225). A total of 20ug of protein was loaded in 1x Laemmli loading buffer with DTT reduction, heated for 2min at 85C and loaded on a TrisGlycine 4-20% SDS-PAGE gel (4-20%, Invitrogen) along with a PrecisionPlus Kaleidoscope protein ladder (BioRad 1610375). Protein was transferred to Immobilon-P PVDF membrane on a TurboBlot station (BioRad). Blots were blocked in 5% BSA block in TBS-T, and stained with the following antibodies in 0.5% BSA in TBS-T: Phospho-TDP-43 (ProteinTech: 22309-1-AP, 1:1000), TDP-43 (Abcam: ab109535, 1:5000), Phospho-Tau (Invitrogen 44752G, 1:1,000), Tau (Invitrogen 13-6400, 1:1000), Amyloid (Biolegend 803004, 1:1000), Vcam (Abcam: ab134047, 1:5000). After two hours with rotation in a sealed bag, blots were washed in TBS-T, and stained with secondard (anti-primary HRP conjugated, at 1:5000) for one hour. They were washed again with TBS-T, and then incubated with luminescent HRP substrate (Fisher PI32109) before imaging on a BioRad ChemiDoc MP system with multiple exposures. Pre-threshold exposures were used for quantitation. Signal was measured in ImageJ, using the line tool and numbers exported to R for data analysis. Signal was normalized to background signal on the blot outside of lane.

## Results

### Erg-based sorting provides a focused analysis of endothelial and microglial alterations across a large set of archived frozen brain tissues

To enrich for endothelial cells (EC) and microglia, cell types typically underrepresented in single nuclei analysis of archived human brain tissues, we tested whether an approach we had applied to isolate these cell types for bulk RNA analysis could also be applied to a droplet based single nuclei analysis approach. We obtained Brodmann Area 10 cortex samples from donors in the NIH NeuroBioBank, including young (N=20, 15-29 years) and old (N=19, 67-95 years) donors without neurodegenerative diseases, or donors with neurodegenerative diseases (N=24 AD, N=20 ALS, N=8 FTD, SI Figure 1 and SI Table 1). Nuclei were prepared and stained as we have previously reported, but with modifications to allow for inCITE-Seq labeling of nuclear proteins, and to obtain a higher yield of sorted nuclei. Consistent with our prior work, we identified clear ERG^Hi^ and ERG^Lo^ populations in the sorted nuclei at approximately the levels we had previously reported, about 2% for Erg^Hi^, 5% for Erg^Lo^ and 20% for NeuN ^17^. The specific ratios varied slightly, but we did not observe obvious trends between disease states (data not shown). Based on our prior work, the ERG^Hi^ nuclei are endothelial, and the ERG^Lo^ nuclei – which likely represent detection of the ERG-like transcription factor FLI1^25^ – are microglia ^17^. Antibody to NeuN, which detects the neuron enriched splice factor Rbfox3, identifies neuronal nuclei ^26^. We sorted nuclei into four groups, endothelial (ERG^Hi^), microglial (ERG^Lo^), neuronal (NeuN), and other parenchymal (ERG^neg^NeuN^neg^), and recombined these to obtain final proportions of 50% endothelial cell nuclei (∼25x enrichment), 25% microglial nuclei (∼5x enrichment), and a mixture of neuronal nuclei and other parenchymal nuclei (Figure 1A). Nuclei were hashtagged, and batches including young, old and two types of dementia samples were included in each 10x reaction. After droplet based 3’RNA analysis of nuclei, alignment of reads, and de-concatenation and removal of doublets by hashtags and Scrublet, we observed proportions very similar to our expected counts, with ∼50% endothelial nuclei overall, and an average of per donor +/− SD (Figure 1B and SI Figure 2). Gene counts and UMI are comparable or higher than previously published datasets (SI Figure 3), and well-known marker genes clearly delineated cell types (Figure 1C). Thus, ERG-based isolation provides a reliable method for the efficient interrogate of endothelial and microglial nuclei across a large set of brain samples.

### Clustering of transcriptional states in sorted nuclei reveals distinct subsets of disease associated expression patterns

Microglial states are well known to correlate with the progression of AD, ALS and FTD ^27–29^. However, less is known about the similarities or differences in endothelial cell transcriptional alterations across neurodegenerative diseases and aging. Existing datasets have examined the question, but typically with either a low number of endothelial cell nuclei across many samples ^15^, or a large number of endothelial nuclei in a few samples ^16^. Here, we leveraged the large number of samples across age and neurodegenerative diseases to determine the consistency in endothelial cell alterations in disease states, and to compare these with healthy aging. An analysis of inCITE seq data revealed well distributed counts of inCITE antibodies between cells (SI Figure 4), and similar clustering of nuclei based on transcript levels to nuclei without inCITE-seq. In general, the quality of the transcript data looked similar, although the mean gene and UMI counts per nuclei were slightly reduced as previously described (SI Figure 4). To identify types of endothelial responses, we clustered capillary endothelial cells from each donor by similarities in gene expression profiles (Figure 2A). This approach yielded three large clusters. The first cluster (1), clearly distinct from the other two, was composed primarily of healthy donors, young and old. Although there were some differences between young donors and older donors within this cluster, these differences were small, relative to the differences between these unaffected individuals and those with any of the neurodegenerative diseases (AD, ALS or FTD). A second cluster (2) was highly enriched for individuals with clinically diagnosed dementia, AD and FTD. A third cluster (3) was composed of older controls and AD, and ALS, with a low portion of FTD samples. This cluster appears to show a deterioration of the pattern associated with cognitively-normal donors, and a partial match to the pattern of expression observed in dementia samples. Although both male and female samples are included, the major differences appeared to be independent of sex. Consistent with the literature, clustering of microglia (Figure 2B) showed specific transcriptional profiles correlated better with cognitively-normal versus dementia samples. In contrast to the endothelium, cognitively normal but older donors generally displayed microglial transcriptional states closer to microglia obtained from donors with dementia. Thus, transcriptional profiles in both endothelial cells and microglia correlate closely with disease progression in AD, ALS and FTD, and surprisingly, the endothelial cell transcriptional patterns associated with dementia-free aging correlate closely with the endothelium of younger donors.

### inCITE-seq identifies disease associated alterations in NF-kB in microglia

Having identified disease associated transcriptional changes in microglia, we aimed to leverage the extensive data on these cells in the progression of AD, FTD and ALS to assess the ability of the inCITE-seq approach to identify alterations in nuclear p65/NF-kB. Increased levels of microglial NF-kB signaling are observed in ALS, FTD and AD, where it contributes to disease progression ^30,31^. Extensive information is now available on microglial states observed by single cell analysis in animal models and human tissues ^29,32^, however there is not yet data linking the microglial phenotypes to specific alterations in p65/NF-kB previously observed in human tissue samples ^33^. To examine NF-kB protein levels in microglial subclusters, we computationally isolated microglial nuclei from all cells, then we clustered inCITE nuclei while regressing out batch effects (from antibody staining and 10x snRNA-seq preparation) but not disease state. This resulted in four clusters of microglia (Figure 3A&B). Analysis of enriched genes identified an enrichment of homeostatic genes in cluster 1 and DAM-associated genes in cluster 0 (Figure 3 &D). Homeostatic genes included CX3CR1 and P2RY12, and DAM associated genes included SPP1 and TMEM163 ^34^. Analysis of inferred pathways implicated NF-kB, TNFα and Trail among the top differential pathways (Figure 3E). Inferred NF-kB activity was highest in cluster 0 associated with DAM markers, and lowest in the homeostatic cluster 1. The enrichment of downstream canonical TNF/NF-kB transcriptional targets was clearly apparent in cluster 0, while these were depleted in cluster 1 (Figure 3F). Analysis of the proportion of each donor that contribute to each of the clusters revealed a significant enrichment of donors from AD, ALS and FTD to DAM-like cluster 0, and a depletion of these same groups in cluster 1 (Figure 3G). Notably, there was a strong correlation of NF-kB Protein levels in Microglia with the level of phospho-Tau measured by Western blot in brain tissue extract from these same samples (Figure 3H), consistent with a proposed positive interaction between these microglia and pTau ^30^. Thus, microglial clustering, and specifically the enrichment of a DAM-like cluster with age, and in AD, ALS and FTD aligns well with expectations based on extensive analysis of these cells in animal models.

We normalized antibody counts to histone H3, as a means of controlling for antibody access to the nuclear compartment. In prior work, nuclear pore proteins have been used for this purpose, but ALS samples exhibited a reduction in nuclear pore proteins consistent with prior studies ^35^. In general, we saw less variation in the levels of H3 measured between disease groups, supporting that this general nuclear marker would be appropriate for normalization (SI Figure 5). We observed an increase in nuclear levels of NF-kB in cluster 0, relative to cluster 1 (Figure 3I). Moreover, when segmenting the antibody staining by relative NF-kB staining intensity (Top 10^th^ percentile, 10^th^-25^th^ percentile and bottom 25^th^ percentile), we observed a clear clustering of lower NF-kB staining in in the homeostatic expressing cluster 1, and highest levels of NF-kB in the DAM associated cluster 0. Notably, the protein data demonstrated a clearer separation of these populations than the inferred NF-kB pathway activity, and delineated an intermediate population of nuclear NF-kB marker by the preferential induction of Trail pathway over the NF-kB pathway (Figure 3E). This was apparent in cluster 2, which appeared to be a transition to the DAM associated cluster 0. An analysis of transcripts positively or negatively associated with TNF/NF-kB signaling revealed a positive correlation of most of these genes in cluster 0 versus other clusters (Figure 3J), and a negative association in cluster 1 (Figure 3K). Examples include IL1B, which is increased in cluster 0 and decreased in cluster 1. Thus, nuclear levels of NF-kB, measured by inCITE-seq, mark a DAM-like cluster with elevated NF-kB transcriptional targets, as well as an intermediate cluster with moderate levels of NF-kB, and activation of TRAIL targets instead of canonical NF-kB targets. This validates the inCITE approach on the well characterized microglial activation in AD, ALS and FTD, and extends previous work by showing that an NF-kB intermediate population (cluster 2) exists in which the pathways activated are not canonical TNF/NF-kB response genes, but more closely resemble TRAIL activation.

### Aging and dementia are indicated by differing nuclear levels of key proteins across subsets of the brain vasculature

Alterations in endothelial cells remain poorly annotated across age and disease states in the human brain, although animal models suggest a prominent role in both vascular aging and neurodegenerative disease ^536^. However, our transcriptional data suggests that alterations within the endothelium are strongly correlated with disease state (Figure 2). Therefore, we performed a similar analysis of brain endothelial cells to the analysis we had performed on microglia. Endothelial cells isolated by ERG^Hi^ included artery, capillary and vein cells, marked by well-described markers (Figure 4A&B). As with microglia, we then subclustered the capillary endothelial cells, and identified 5 major clusters (Figure 4C). Analysis of these clusters revealed cluster 3 to be almost entirely composed of cells from cognitively-normal donors, young and old (Figure 4D&E). On the other hand, cluster 1 was composed almost entirely of cell from donors with neurodegenerative disease (Figure 4D&E). The age of the donor had little effect on the contribution of cells to these clusters, as both young donors and cognitively-normal aged donors exhibited a heavy skewing towards cluster 0 and away from cluster 1. Together, these two clusters represent approximately 50% of all of the capillary endothelial cells from each donor (Figure 4E), and are therefore a large portion of the capillary bed. Analysis of pathways enriched in the disease associated cluster 1 revealed increased expression of pathways including epithelial-mesenchymal transition, G2M checkpoint, and TNF/NF-kB signaling (Figure 4F). Thus, analysis of endothelial populations across 89 donors reveals a switch in endothelial cell states in neurodegenerative diseases AD, ALS and FTD, which are highly enriched in cluster 1 versus cluster 3.

### inCITE-Seq analysis shows a reduction in nuclear β-Catenin and Wnt target genes in the disease enriched capillary endothelial cell cluster

The inclusion of inCITE-seq antibodies allowed us to take this analysis one step further, and to assess key markers of endothelial activation (p65/NF-kB) and barrier function (β-Catenin/Wnt) and TDP-43 across these clusters. Analysis of antibody staining intensity across cell types showed an increase in nuclear β-Catenin in healthy cluster 3 and a decrease is disease cluster 1, a similar trend to TDP-43 (Figure 5A&B). Density plot analysis, showing the frequency of cells with the top 10^th^ percentile of β-Catenin, showed a clear reduction in cluster 1 versus cluster 3 (Figure 5C). This coincided with an enrichment of upregulated transcriptional markers of β-Catenin activation in cluster 3 versus cluster 1 (Figure 5C&D). Key transcriptional targets TCF/LEF1, ABCG2 and APCDD1 were enriched in cluster 3 versus cluster 1 (Figure 5E&F). Thus, the cluster 1 population, which is highly enriched for AD, ALS and FTD donors, exhibits reduced levels of TDP-43 and nuclear β-Catenin, and reduced expression of canonical Wnt response genes.

### Reduced nuclear levels of TDP-43 define a specific transcriptional state unique to healthy aging and not observed in AD and ALS/FTD

There was a small, but statistically significant increase in p65/NF-kB in healthy cluster 3 versus disease cluster 1 (Figure 5A). However, there was a prominent and opposite increase in NF-kB transcriptional targets in disease cluster 1 versus healthy cluster 3 (Figure 5A). To shed light on this paradox, we obtained insight from our parallel studies of TDP-43 function in animal models and cultured cells (BioRxiv reference). This work indicated that a reduction in TDP-43 could lead to increased activation of p65/NF-kB transcriptional targets in brain ECs at steady state. Thus, the reduction in TDP-43 we had observed in the nuclei in this disease associated capillary cluster might regulate the transcriptional response associated with nuclear p65/NF-kB. Therefore, we examined the level of nuclear TDP-43 together with p65/NF-kB by our inCITE-seq approach.

As TDP-43 lacks the well-defined transcriptional output of NF-kB and Wnt, we validated our inCITE-seq staining by examining cells with siRNA-mediated suppression of TDP-43. We observed in substantial reduction in detected TDP-43 with siRNA treatment in a human brain endothelial cell line, HBEC5i (Log2-FC, Pval, SI Figure 6A). Furthermore, as nuclear levels of TDP-43 are reduced in >90% of cases of ALS and >50% of AD, we examined nuclear levels of TDP-43 in neuronal nuclei from ALS and AD donors. We found a reduction in these nuclei as well (SI Figure 6B). Thus, our inCITE-measurement of TDP-43 accurately reflects experimental and expected loss of nuclear TDP-43.

We then examined how the levels of nuclear TDP-43 changed across the clusters. In particular, we focused on the cells with the highest level of nuclear p65/NF-kB, which we believed would be driving the transcriptional NF-kB response detected in our data. We found that, although there were high levels of nuclear p65/NF-kB in both cluster 1 and cluster 3, this was associated with a reduction in TDP-43 specifically in the AD, ALS and FTD enriched cluster 1 (Figure 6A). We also noted that the distribution in the density plot was very different, with p65/NF-kB high cells in disease cluster 1 being tightly packed, but distributed in cluster 3. As UMAP clustering proximity reflects similarities in gene expression patterns in each nucleus, this suggests a strong and conserved transcriptional signature across cells with high levels of nuclear p65/NF-kB in diseased endothelial cells in cluster 1, but not in healthy endothelial cells in cluster 3. In line with this observation, we observed strong alterations in gene expression correlated with increasing levels of p65/NF-kB in disease cluster 1, but not healthy cluster 3 (Figure 6B). Notably, the specific transcripts and pathways associated in each were also different, and pathways associated with p65/NF-kB levels in diseased endothelial cells included EndoMT, TGFβ and canonical TNF/NF-kB signaling, while the pathways associated with NF-kB in healthy endothelial cells included interferon, JAK/STAT and apical junction (Figure 6C).

To examine the relationship between TDP-43 and NF-kB more closely by disease state, we plotted the relationship of NF-kB to TDP-43 levels in each samples type, hypothesizing that the stoichiometry may need to be maintained between their nuclear levels. On these plots, a continuum of cells can be observed with a proportional increase in TDP-43 with NF-kB (Figure 6D). This relationship is almost identical in cognitively normal donors, young or old, and AD, ALS and FTD donors, until NF-kB levels rise to the highest levels (Top 25^th^ percentile, Figure 6D). At this point, the relationship breaks down, most prominently in ALS, but also, to a lesser extent, in AD and FTD, and the levels of TDP-43, relative to NF-kB, drop (Figure 6D, with quantitation in Figure 6E). Importantly, these subsets included a large number of nuclei from several donors, indicating similar responses across the disease group (SI Figure 7).

**Figure 7.**
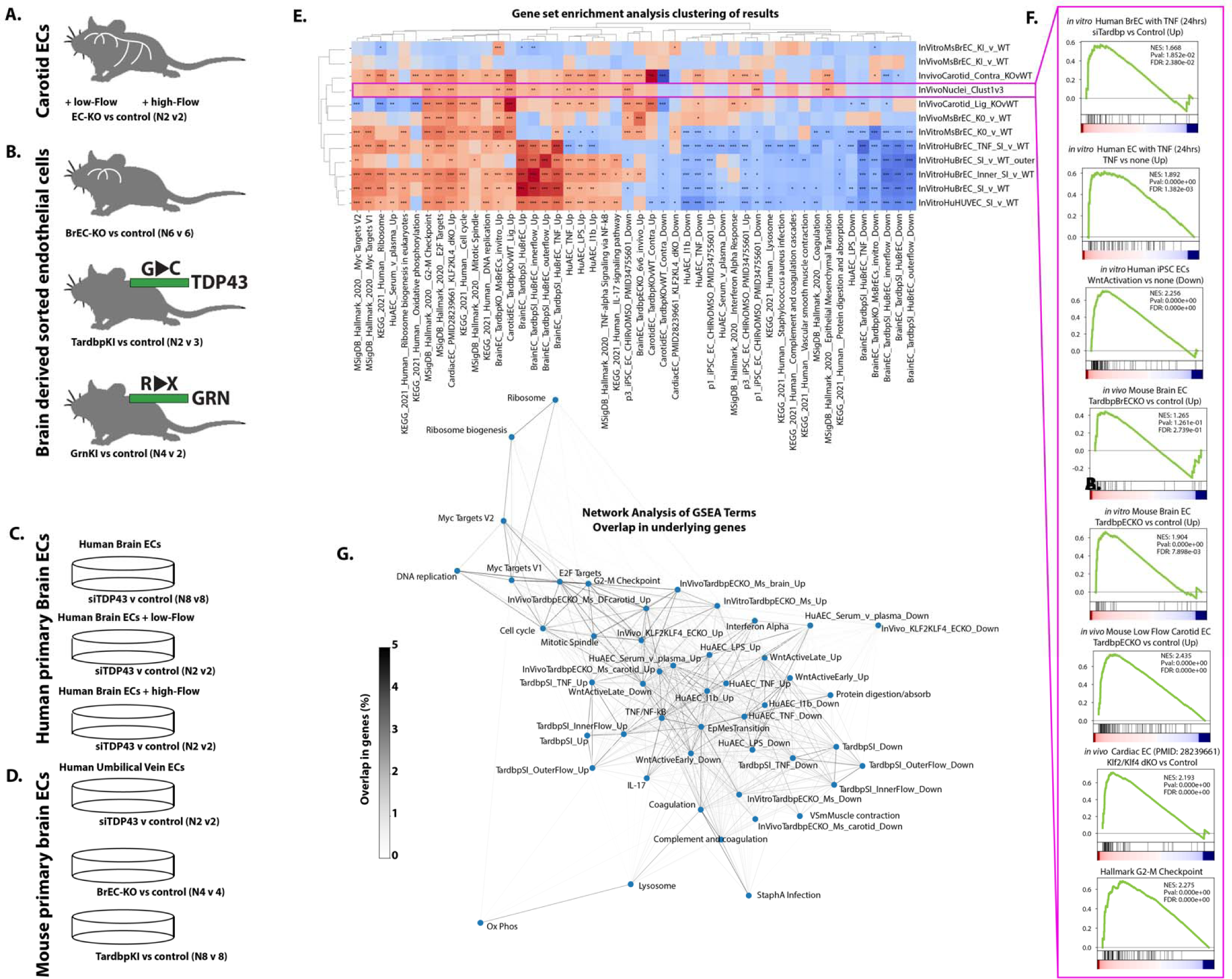
Transcriptional alterations associated with experimental loss of TDP43. (A-D) Source data and replicate numbers for differential expression analysis. (E) Heatmap showing the normalized enrichment score (NES) for specific KEGG and Hallmark pathways and custom gene sets for endothelial cell pathways by GSEA analysis, only pathways achieving an FDR<0.01 in two or more comparisons are shown. (F) Example GSEA enrichment plots, showing pathway enrichment in disease versus healthy capillary clusters in cortex (Cluster 1 vs Cluster 3). (G) Clustering of GSEA pathways enriched by overlap in genes between pathways. (E) GSEA derived *FDR<0.05, ** FDR<0.01, ***FDR<0.001.

Together, our data shows that nuclear levels of TDP-43 are reduced specifically in AD, ALS and FTD donors at the highest levels of NF-kB, and that this correlates with an increased level of TNF/NF-kB target genes in these cells.

### Transcriptional alterations in TDP-43 depleted endothelial cells correlates with experimental depletion of nuclear TDP-43

Our data from human samples is correlative, but suggests the possibility that a loss in nuclear TDP-43 alters the cellular response to nuclear p65/NF-kB levels. However, alterations in TDP-43 are difficult to isolate from the many other changes in AD, ALS and FTD, and assignment of cause is difficult. To directly assess the consequence of the loss of nuclear TDP-43 in the brain endothelium, we performed experiments across mouse models of ALS/FTD with point mutations in TDP-43 or GRN, leading to reduced nuclear levels of TDP-43, as well as a targeted mouse model of brain endothelial TDP-43 deletion, and cultured mouse and human primary brain endothelial cells with these same mutations or silencing of TDP-43 (Figure 7A-D, BioRxiv reference). Assessing pathway enrichment across all of these conditions by GSEA analysis, we observed that the targeted loss of TDP-43 led to responses consistent with the difference in transcription between TDP-43 high cluster 3 and TDP-43 low cluster 1 (Figure 7E). Including other published data, and our own data on the treatment of endothelial cells with a range of stimuli (e.g. proinflammatory cytokines, disturbed flow, and serum or plasma), we observed a very similar transcriptional signature in human primary brain endothelial cells treated with TNF, or in the presence of disturbed flow, and in mouse brain or arterial endothelial cells with targeting of a floxed Tardbp/TDP-43 gene (Figure 7E&F). Other pathways notably altered in both cells with targeted disruption of TDP-43 and disease cluster 1 include Wnt (downregulated), and genes increased at G2-M checkpoint and with deletion of key endothelial maintenance transcription factors KLF2 and KLF4 (Figure 7E&F). Network analysis of the genes found within each of these pathways showed that, while the signatures for each gene module were distinct (typically <1-2% overlap) that they formed clusters, with pro-inflammatory genes (e.g. TNF and IL1b response in human aortic ECs) being related to Wnt regulated and KLF2/KLF4 regulated genes and distinct from ribosome, and myc-target genes.

Thus, an analysis of murine and human endothelial cells with partial and complete endothelial loss of TDP-43 also identifies increased NF-kB target genes and decreased Wnt target genes, and extends this to show several other signaling pathways consistently affected in cells with the targeted loss of TDP-43 and the cluster 1 ECs with reduced levels TDP-43. Together, this implicates the loss of TDP-43 occurring in cluster 1 capillary ECs as a driver of the increased NF-kB transcriptional responses and decreased Wnt/β-Catenin signaling in these disease associated ECs.

## Discussion

Here, we show that nearly half of the capillary endothelial cells across a range of neurodegenerative disease states, including AD, ALS and FTD, exhibit a unique transcriptional signature. By applying inCITE-seq analysis of TDP-43 and key transcription factors, we reveal that an increased canonical NF-kB transcriptional output in disease associated endothelial cells is not associated with increased levels of nuclear NF-kB, but is instead associated with a drop in nuclear levels of TDP-43 at the highest levels of NF-kB. This correlates with increased expression of transcripts involved in immune cell recruitment and coagulation, and a reduction in a key regulator of endothelial cell barrier function, Wnt signaling. Changes in the levels of TDP-43 in the endothelium are sufficient to drive many of the transcriptional alterations observed in disease population capillary endothelial cells, implicating a disease-specific loss of nuclear TDP-43 from endothelial cells in the loss of BBB integrity observed in AD, ALS and FTD.

### Impaired coordination of NF-kB and TDP-43 in neurodegenerative capillary endothelial cells

The discovery of elevated nuclear p65/NF-kB levels in both cognitively normal and neurodegenerative brain endothelial cells highlights a paradox. In epithelial cells and endothelial cells, NF-kB appears to be critical the barrier maintenance, a role diametrically opposed to its canonical function in inflammatory responses and barrier disruption. In endothelial cells, nuclear translocation and transcriptional activation by NF-kB forms the core of the inflammatory response to pathogens and cytokines, culminating in increased leukocyte recruitment and permeability ^37^. However, it is also apparent that some basal level of NF-kB signaling is required to maintain tight barriers. Deletion of Nemo, a key component in NF-kB signaling, results in increased intestinal barrier permeability ^38^, and NF-kB inhibitors disrupted tight junctions and increased lung epithelial cell permeability doses which did not induce cell death ^39^. A similar role appears to exist in brain endothelial cells, where loss of a critical components of the NF-kB signaling pathway, Nemo and Tak1, lead to BBB permeability, endothelial cell death and vascular rarefication ^40^. Consistent with our results, high levels of basal p65/NF-kB were also observed murine brain endothelial nuclei ^18^. However, these high basal levels of nuclear p65/NF-kB somehow appear not to activate canonical downstream transcriptional responses, which are low in the murine brain endothelium ^41^ and in our isolated human endothelial nuclei. The molecular mechanisms underlying this paradox have been unclear.

Our research sheds light on this paradox, particularly regarding the significant impact of TDP-43 disruption on brain endothelial cells. Comparative analysis of healthy and diseased endothelial nuclei reveals a unique pattern in the diseased group: although the spectrum of nuclear p65/NF-kB levels is comparable in both groups, only the diseased group demonstrates a pronounced canonical NF-kB response. As this correlates with a drop in the levels of nuclear TDP-43 relative to NF-kB, we propose that TDP-43 limits NF-kB-mediated activation of canonical transcriptional responses while allowing non-canonical functions in barrier maintenance. Our data from targeted disruption of TDP-43, *in vivo* and *in vitro*, support this concept. We observe increased levels of canonical TNF/NF-kB targets in endothelial cells with deletion or suppression of TDP-43. Thus, we propose that TDP-43 levels rise with nuclear levels of NF-kB, and that this relationship must be maintained to allow NF-kB to maintaining the BBB without triggering transcriptional activation of well-known proinflammatory pathways. Our data show that this relationship is disrupted in AD, ALS and FTD.

The molecular mechanisms underlying this relationship remain unclear. TDP-43 directly interacts with NF-kB, but this relationship, at least in microglia, has been shown to increase activation of a canonical transcriptional target – opposite to our suggested model ^42^. Some data has also suggested that TDP-43 may limit nuclear translocation of p65/NF-kB in a cell-type dependent manner ^43^. However, this also does seem to explain the inhibition we observe, as different transcriptional responses are observed at the same levels of nuclear p65/NF-kB. Other possible mechanisms, including sequestration of p65/NF-kB in the nucleus, and TDP-43-mediated regulation of co-factors and competition at canonical transcriptional binding sites should be explored.

### Identification of a disease associated capillary endothelial cluster associated with reduced Wnt/β-Catenin

The Wnt signaling pathway is a critical mediator of blood brain barrier formation and maintenance, however its alterations in the progressive barrier dysfunction in neurodegenerative disease has not been addressed in human tissues ^44^. Although some changes in Wnt ligand and negative regulators have been observed in the brain tissue in AD, is not clear whether this leads to altered Wnt signaling in some or all endothelial cells. Here, using both a protein-based readout of nuclear β-Catenin and a transcriptional readout of Wnt signaling activity in the same cells, we show a reduction in both measures in a subset of brain endothelial cells associated with AD, ALS and FTD. Although it is not clear why Wnt signaling levels are reduced in these neurodegenerative disease states, an intriguing possibility is that it is linked to TDP-43 loss. While the overlapping loss of TDP-43 and Wnt in capillary cluster 1 is correlational, data from cells with targeted deletion or suppression of TDP-43 *in vitro* or *in vivo* showed a similar increase in Wnt suppressed transcripts and decrease in Wnt induced transcripts. In addition, loss of TDP-43 in primary human brain endothelial cells lead to altered splicing of key Wnt regulators DKK1 and β-Catenin itself, CTNNB1. Notably, in a companion paper, we show that even small perturbations of TDP-43 (e.g. ALS/FTD associated heterozygous mutation) lead to significant barrier defects in mice harboring a single heterozygous ALS/FTD associated mutation in the gene. This might suggest that the loss of TDP-43 and loss of Wnt/ β-Catenin in diseased cluster 1 are driven by a common mechanism, namely the reduction in nuclear levels of TDP-43. Future work, focused specifically on these signaling pathways in tractable animal models will be required to delineate these interactions.

In summary, by applying a novel method for endothelial and microglial enrichment for relatively high throughput analysis of a large set of human cortical brain tissues across age and neurodegenerative disease states, we validate inCITE-seq as a powerful approach to dissecting signaling pathways by the measurement of nuclear transcription factors and RNA-binding proteins. Using this approach, we define a previously unappreciated relationship between p65/NF-kB and TDP-43 in the nuclei of brain capillary endothelial cells, which determines the type of transcriptional output that occurs, and defines a subtype of pro-inflammatory and barrier compromised endothelium unique to a range of neurodegenerative diseases, including AD, ALS and FTD. A better understanding of the molecular regulation in these endothelial cells is likely to provide improved biomarkers for early vascular alterations in the progression of these diseases, and avenues to limit barrier and vascular dysfunction contributing to their progression.

## Acknowledgements

This work was supported by UConn Health startup funds from the School of Medicine and Department of Cell Biology, Center for Vascular Biology and Calhoun Cardiology Center, American Heart Association Innovative Project Award 19IPLOI34770151 (to P.A.M.); NIH National Heart, Lung, and Blood Institute Grants K99/R00-HL125727 and RF1-NS117449 (to P.A.M); American Heart Association Predoctoral award (to O.M.F.O.) and NIH GM135592 (to B.H.). We are grateful for the consistent help from Vijendar Singh and the staff at the Computational Biology Core at UConn Health in providing input on computational approaches and installing packages on the cluster. We are appreciative of the assistance of the staff at the NIH NeuroBioBank in preparing tissue samples and detailed reports, and for the donors who provided tissues to advance knowledge. Finally, we appreciate the help of the Bernard Cook, Science Writer and Illustrator in the Dean’s Office at the School of Medicine at UConn Health, who prepared illustrations for this work, and for other members of the lab, who provided consistent input into the research described here.

## Author contributions

Omar M.F. Omar (OMFO), Amy L. Kimble (ALK), Ashok Cheemala (AC), Jordan D. Tyburski (JDT), Swati Pandey (SP), Qian Wu (QW), Bo Reese (BR), Evan R. Jellison (ERJ), Yunfeng Li (YL), Bing Hao (BH), Riqiang Yan (RY), Patrick A. Murphy (PAM)

PAM and OMFO were responsible for conceptualizing the study. OMFO conducted the antibody conjugation. OMFO performed single nuclei experiments with help from JDT and ERJ. OMFO performed computational analysis of resulting data with input from PAM. QW consulted on sample selection. YL and BH provided expertise and conducted the antibody purification using size exclusion chromatography. ALK and BR performed library preparation and sequencing. SP performed comparative analysis with other single nuclei data sets. AC provided the cell line necessary for the knockdown experiment, and data from Tardbp siRNA in human cells and Tardbp KO and KI in mouse cells *in vitro* and *in vivo*. JDT performed Western blot analysis. The manuscript was primarily written by PAM and OMFO, with input all authors, including RY, contributing through review and input.

---

**SI Figure 1.**
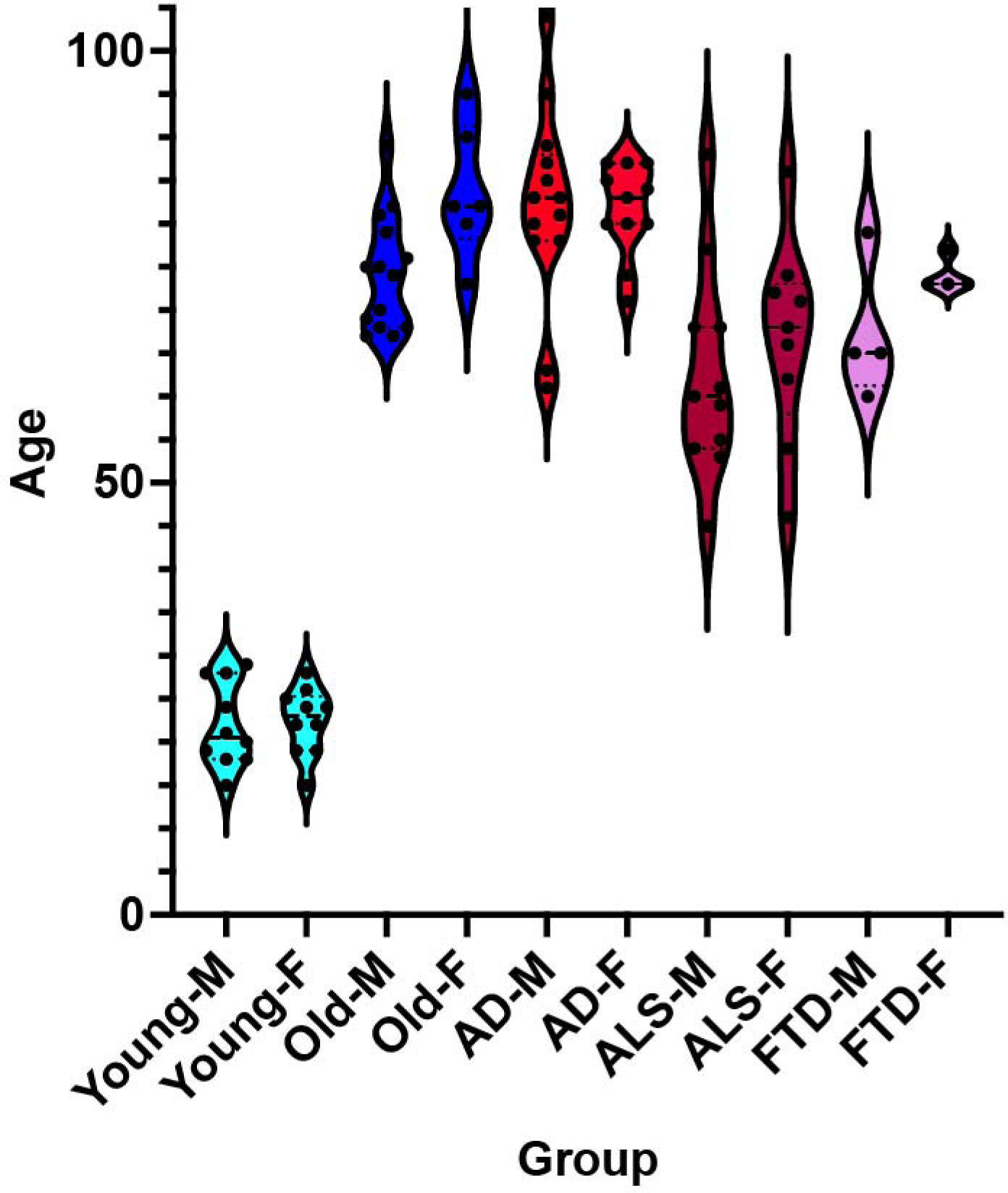
Age and sex distribution of cortical samples used. Plots show the age of individual sample and their distribution.

**SI Figure 2.**
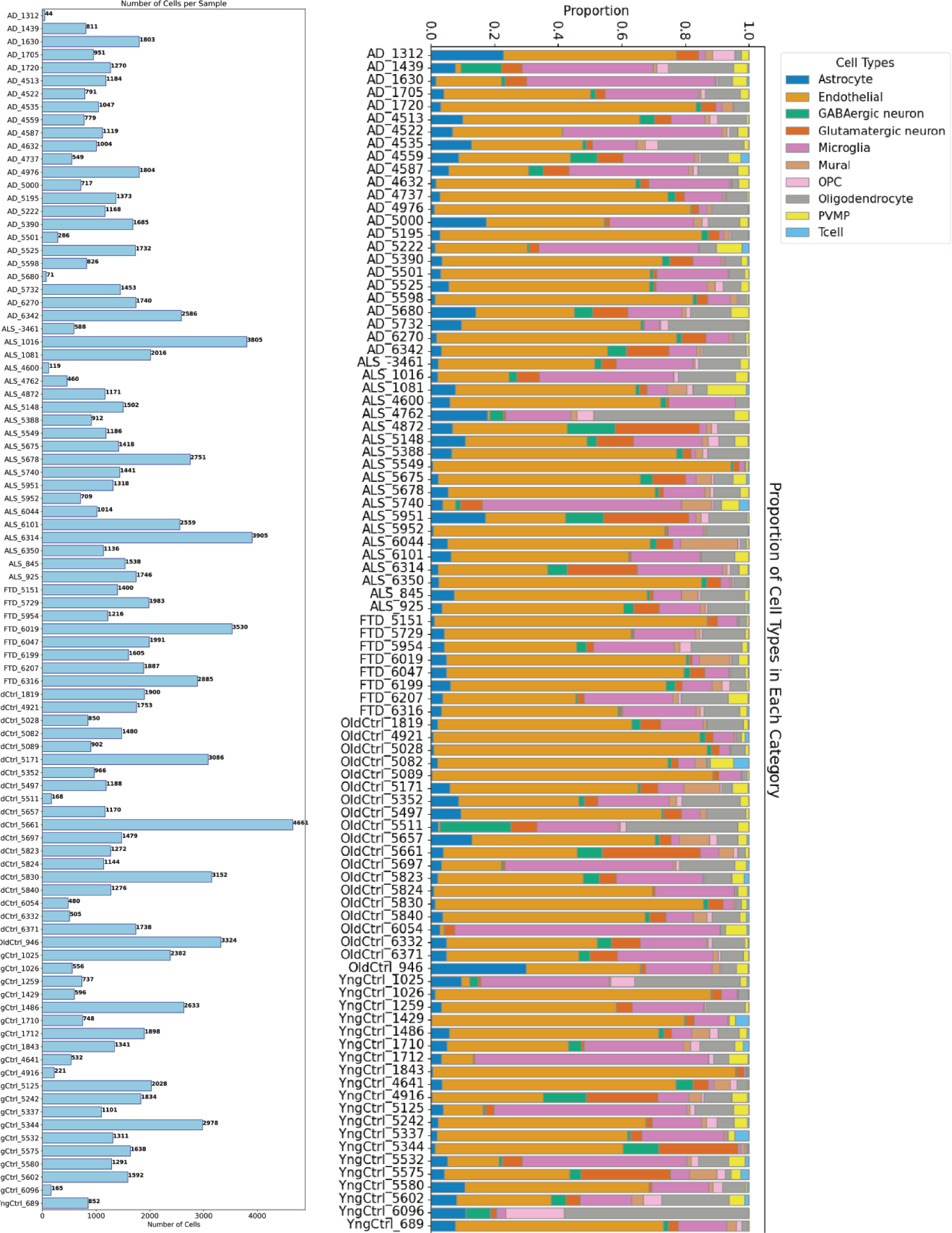
Number of nuclei and distribution of cell types in data. Sample ids are shown on the y-axis, and plots show total nuclei (with count), endothelial nuclei (with count), and distribution of cell types (key in top left).

**SI Figure 3.**
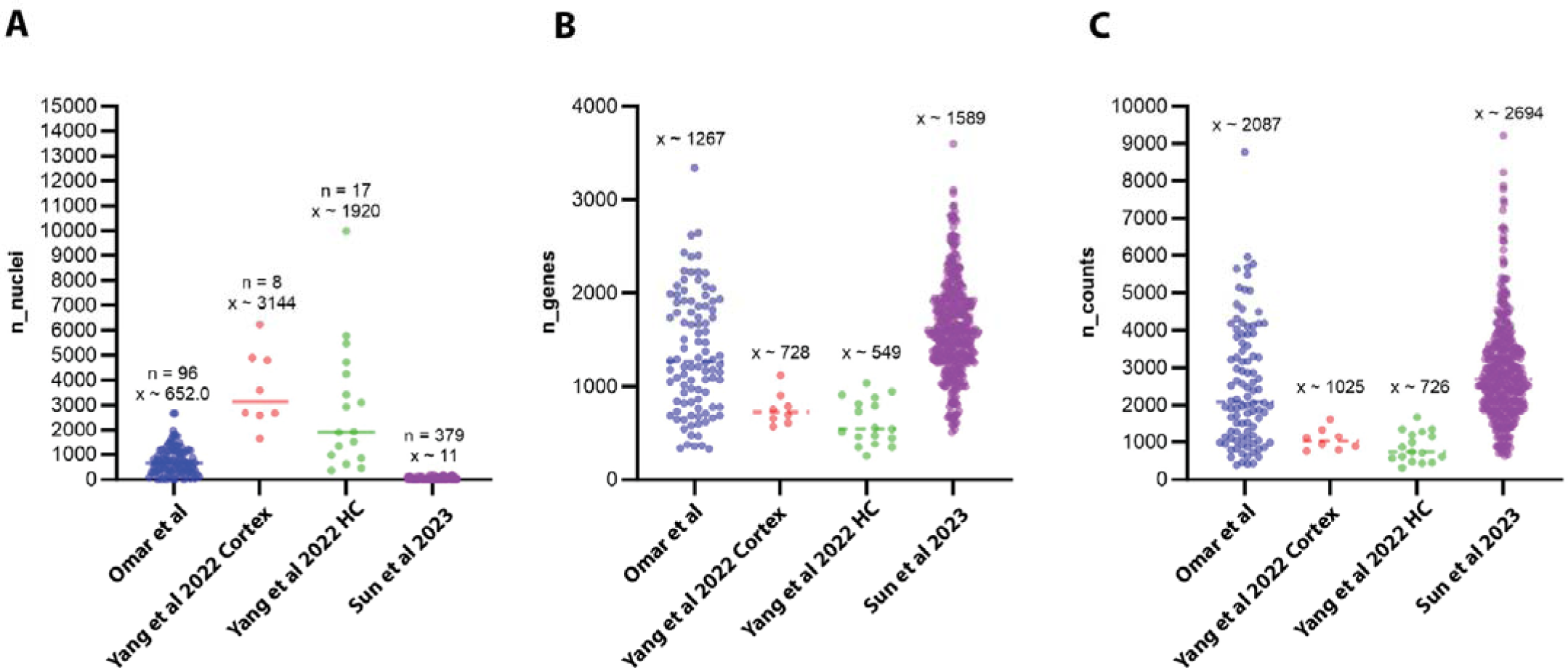
Comparison of gene counts and UMI with prior data sets. Comparison of number of nuclei per donor, and number of genes and counts per donor in data described here, relative to recently published data sets on brain vasculature and endothelium.

**SI Figure 4.**
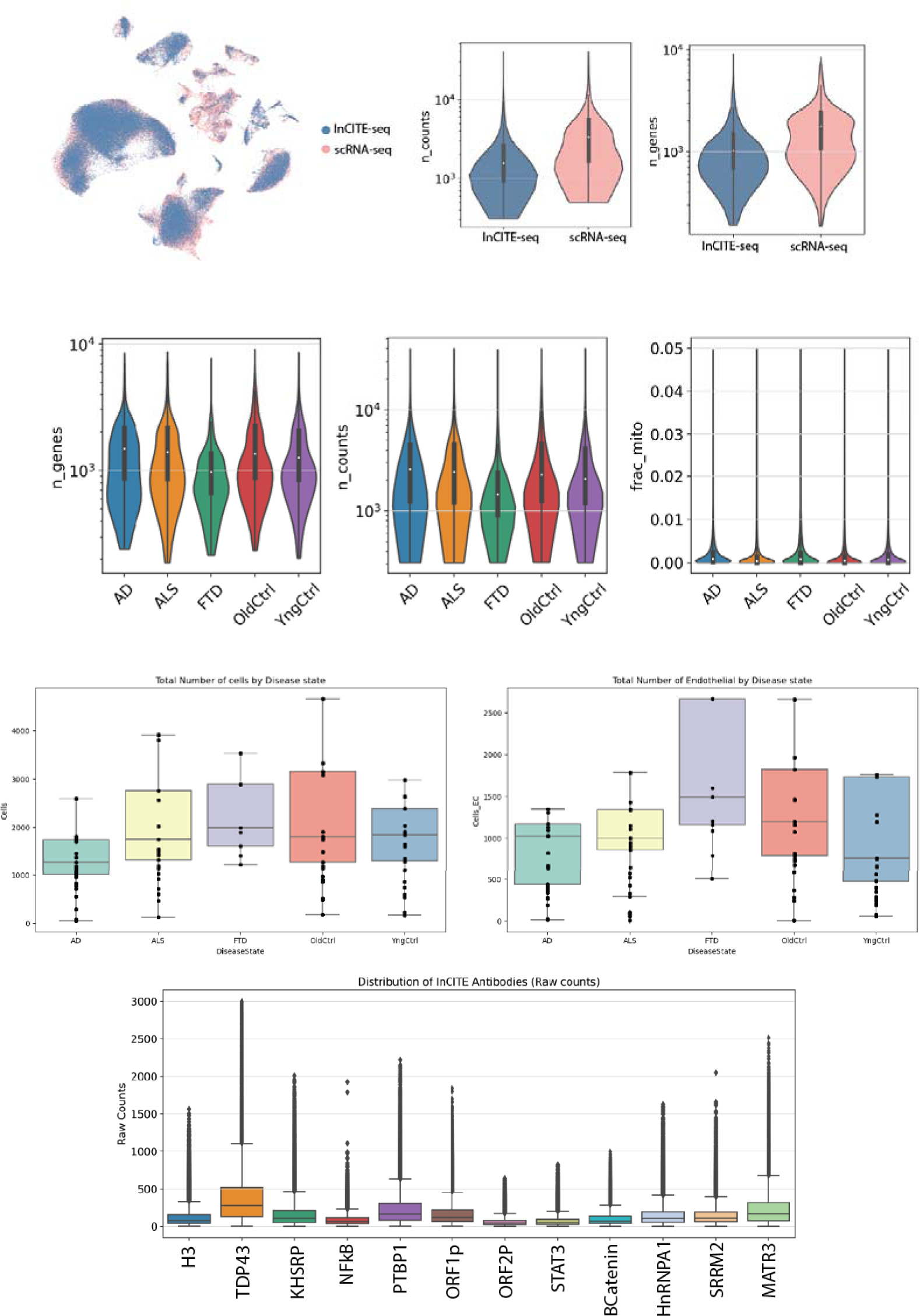
Comparative gene counts and umi in single nuclei data (with and without inCITE-Seq) UMAP showing overlap between single nuclei data obtained from standard scRNA-Seq and inCITE-Seq preparations. UMI counts per cells and gene counts per cell are shown overlap, and by subcategory, along with mitochondrial RNA counts. Plots show the total number of cells by disease state in each method.

**SI Figure 5.**
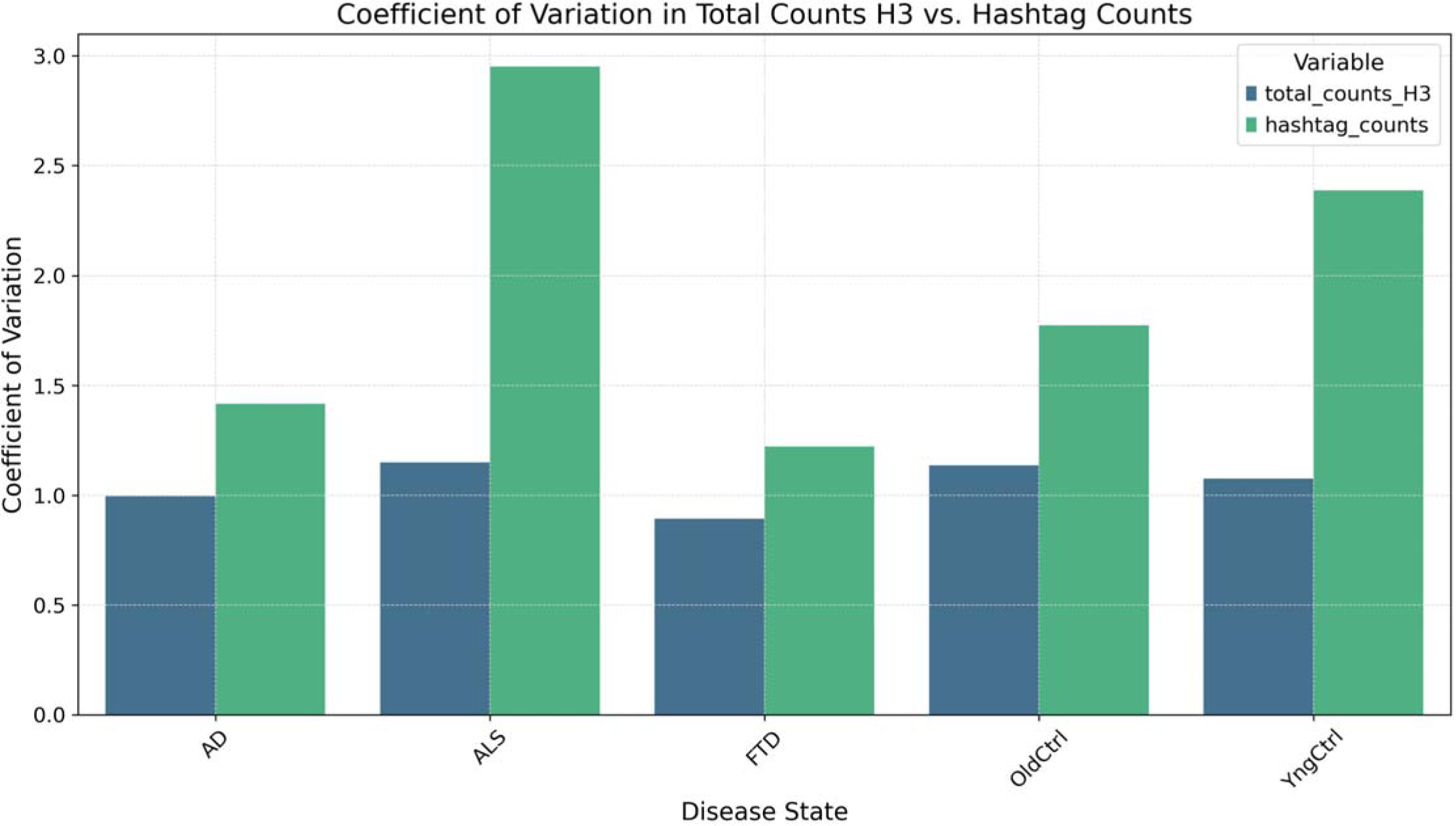
Coefficient of variation in the levels of histone and nuclear pore hashtags between nuclei by disease state. Violin plot shows coefficient of variation in levels of histone and nuclear pore hashtags across nuclei within each disease group.

**SI Figure 6.**
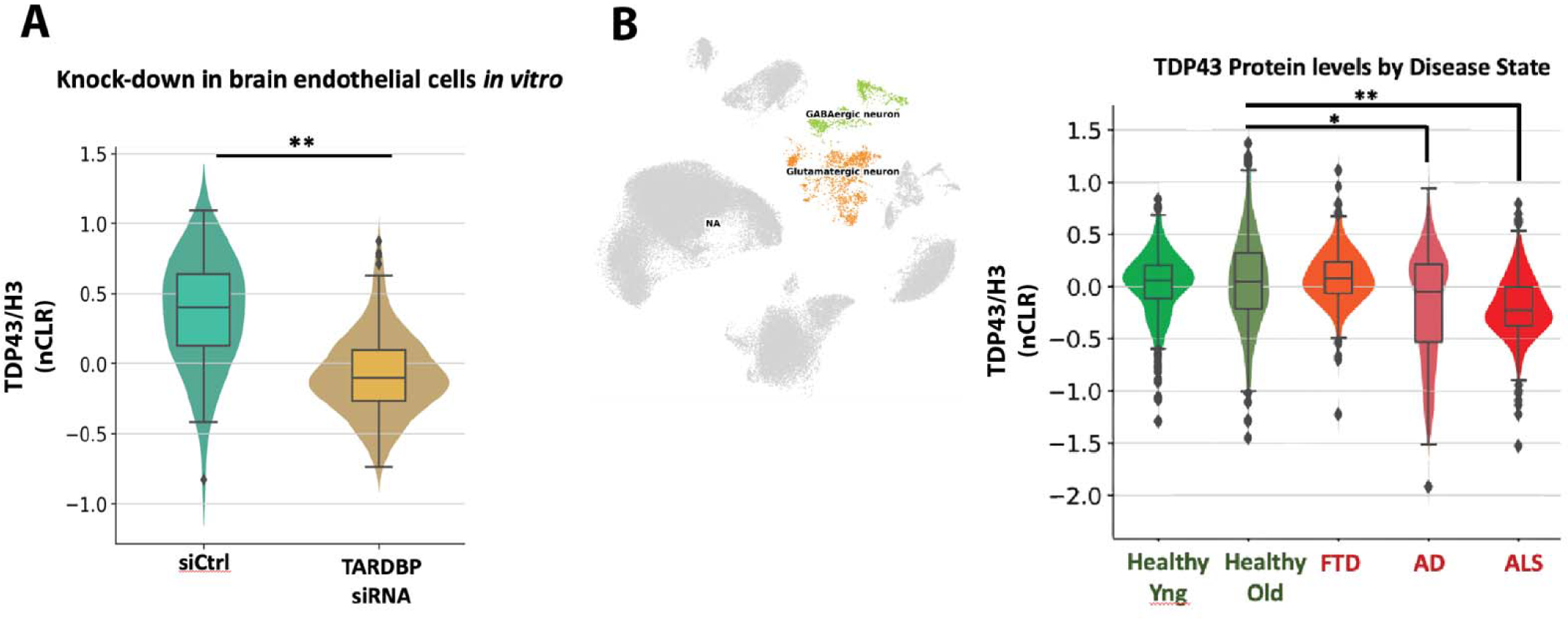
Validation of TDP-43 inCITE analysis. (A) siTDP-43 knockdown in HBEC5i human brain endothelial cell line, compared to siControl treated HBEC5i cells, and analyzed in an inCITEseq experiment along with a standard brain nuclei preparation. (B) UMAP showing neuronal clusters analyzed, and violin plots showing the levels of nuclear TDP-43 protein (relative to nuclear H3) by disease state in these neuronal clusters.

**SI Figure 7.**
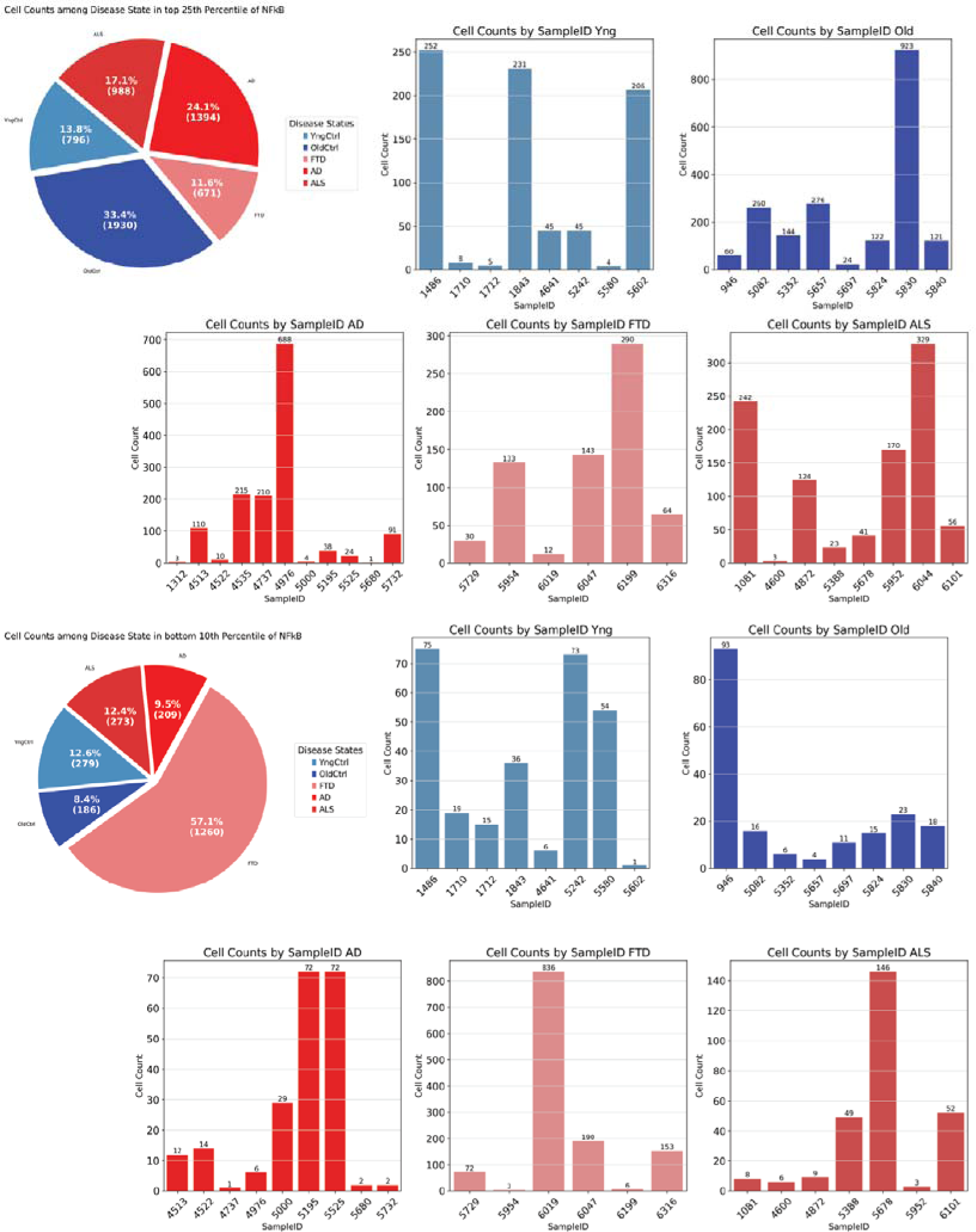
Composition of samples in extremes of p65/NFkB. Pies chart bar plots showing the proportion of cells in the top 25^th^ percentile of p65/NFkB:H3 and the bottom 10^th^ percentile of p65/NFkB:H3.

**SI Figure 8.**
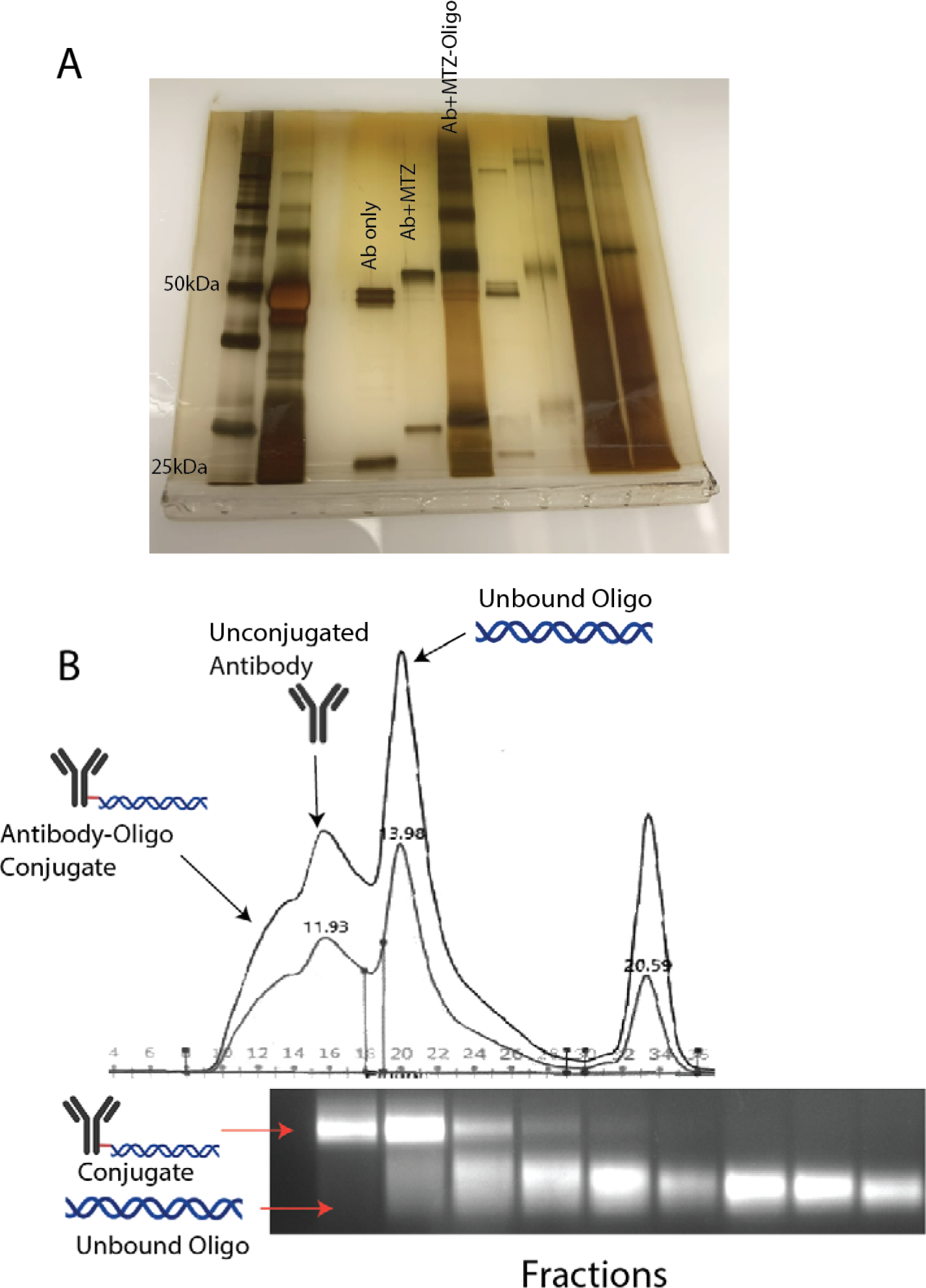
Size-exclusion purification of oligo conjugated antibodies. (A) Reduced and silver stained gel showing the step wise addition of click chemistry adduct (MTZ) and oligo nucleotide (MTZ-Oligo). (B) Oligo-conjugated antibody cleanup via size exclusion chromatography. Top, a representative size exclusion chromatogram recorded with dual absorption wavelengths (260 and 280 nm). Conjugated antibodies, having the largest molecular weights, were eluted first from the column, followed by the free antibodies. Unconjugated oligonucleotides were the last to be eluted from the column. Bottom, fractions from the column were analyzed by an agarose gel stained with SybrGold showing the presence of free and conjugated oligonucleotides.

## Notes

### Competing Interest Statement

The authors have declared no competing interest.

